# A case for glycerol as an acceptable additive for single particle cryoEM samples

**DOI:** 10.1101/2021.09.10.459874

**Authors:** Benjamin Basanta, Marscha M. Hirschi, Danielle A. Grotjahn, Gabriel C. Lander

## Abstract

Buffer composition and sample preparation guidelines for cryo-electron microscopy are geared toward maximizing imaging contrast and reducing electron beam-induced motion. These pursuits often involve the minimization or complete removal of additives that are commonly used to facilitate proper protein folding and minimize aggregation. Among these admonished additives is glycerol, a widely used osmolyte that aids protein stability. In this work, we show that inclusion of glycerol does not preclude high-resolution structure determination by cryoEM, as demonstrated by a ∼2.3 Å reconstruction of mouse apoferritin (∼500 kDa) and a ∼3.3 Å reconstruction of rabbit muscle aldolase (∼160 kDa) in presence of 20% v/v glycerol. While we found that generating thin ice that is amenable for high-resolution imaging requires long blot times, the addition of glycerol did not result in increased beam-induced motion nor an inability to pick particles. Overall, our findings indicate glycerol should not be discounted as a cryoEM sample buffer additive, particularly for large, fragile complexes that are prone to disassembly or aggregation upon its removal.

## Introduction

The initial, and often most challenging, task associated with detailed structural characterization of a biological target is the biochemical purification of the macromolecule, and this remains a significant bottleneck in the field of structural biology. Often, substantial efforts are placed on optimizing the recombinant expression or reconstitution of the targeted macromolecule and subsequent isolation of the target into a solution that optimally preserves its structural composition and stability. Glycerol is a linear, three-carbon polyol widely used in buffer solutions to enhance macromolecular stability, since glycerol prevents protein unfolding by stabilizing aggregation-prone intermediates and favoring more compact protein conformations (Vagenende *et al*., 2009). Many purified complexes strictly require the presence of glycerol for biochemical stability and are susceptible to unfolding or aggregation in its absence. Despite the notable benefits for target stability, the inclusion of glycerol in the final purification buffer has been strongly discouraged by the cryo electron microscopy (cryoEM) community.

There are three main arguments that have been made against the inclusion of glycerol in cryoEM buffers: First, the inclusion of glycerol in the buffer increases the solvent density which is thought to decrease contrast to the detriment of data quality and structure solution. The density of a 20% glycerol solution at -200 °C is 1.033 g/cm^3^ while the average density of protein is 1.181 g/cm^3^. Given that our ability to resolve macromolecular structures embedded in vitrified buffer depends on the differential scattering of electrons of the target complex vs. the surrounding buffer, it is thought that increasing the density of the buffer will decrease attainable image contrast. Second, glycerol was previously shown to significantly increase beam-induced motion during image acquisition. For instance, addition of 50% glycerol increased the motion of gold fiducials up to 10-fold (Karuppasamy *et al*., 2011). Motion of the sample during acquisition will blur highresolution features, and even motion correction methodologies cannot recover high resolution information from images containing substantial beam-induced motion. Third, it is widely believed that addition of glycerol to the vitrification buffer increases sensitivity to radiation damage, as “bubbling” is often observed in EM samples containing glycerol (Frederik *et al*., 1991, Karuppasamy *et al*., 2011). This bubbling is thought to be caused by the production of hydrogen gas upon irradiation of biological materials and small organic molecules such as glycerol. Thus, the presence of glycerol in the buffer may cause widespread bubbling and encumber structure determination.

These arguments against the use of glycerol in sample preparation for cryoEM structure determination resulted in widespread avoidance of glycerol, even when macromolecular complexes require the biochemical stabilization it provides. However, the limited literature on the subject describes experiments involving glycerol concentrations of 50% v/v (Karuppasamy *et al*., 2011), which is higher than those typically considered for aiding solubility during protein purification (5-40% v/v) (Bondos & Bicknell, 2003), imaging thick, dense biological samples that bear little resemblance to those prepared for single-particle analysis (Frederik *et al*., 1991), and use relatively low imaging voltages (100-120keV), which are known to cause more radiation damage than the current voltages typically used for single particle cryoEM (Peet *et al*., 2019). In order to expand the utility of cryoEM imaging for samples that require glycerol for stability, we tested the effects of a buffer containing 20% glycerol on our ability to solve high resolution reconstructions of two model specimens, apoferritin and aldolase, using current imaging methodologies.

The chosen glycerol concentration is the midpoint between the maximum generally considered for protein stabilization during purification (40% v/v), and the concentration deemed practical for size exclusion chromatography (*<*10% v/v), a ubiquitous step in protein purification. As expected, the increased viscosity of the buffer made optimal grid preparation more challenging and the associated increase in ice thickness complicated image analyses. We were nonetheless able to determine high resolution structures of both model specimens. These findings demonstrate that the inclusion of glycerol in the final cryoEM buffer of a targeted sample does not abolish the ability to determine a high-resolution structure, thereby widening the range of biological targets that can be studied with cryoEM.

## Results

### High resolution structure determination of apoferritin in presence of 20% glycerol imaged at 200 keV

In order to determine the effect glycerol has on the resolutions of structures that can be obtained using single particle analysis, we acquired data on mouse heavy chain apoferritin in the presence of 1.66% and 20% (v/v) glycerol using a 200 keV Talos Arctica. The apoferritin protein assembles into a ∼500 kDa octahedron, and is widely used for cryoEM studies due to its size and stability, and has been resolved to better than 2 Å resolution in prior studies (Yip *et al*., 2020). Notably, inclusion of 20% glycerol did not impede our ability to determine high-resolution structures, as both datasets (+/-glycerol) gave rise to reconstructions of apoferritin with reported resolutions at the Nyquist frequency (2.3 Å) (Fig. 1A and Supplementary Fig. 7D). Moreover, the densities of both reconstructions show a tight local resolution distribution near 2.3 Å, not surpassing 2.4 Å (Fig. 1B and Supplementary Fig. 7F), and the density quality and detail of local features are indistinguishable between the two reconstructions (Fig. 1C). However, the higher glycerol concentration indeed appeared to lower the overall quality of the data, as more extensive processing was involved in attaining the final structure (Supplementary Fig. 1 and 2), a greater number of particles was required to achieve high resolution (consistent with the higher B-factor shown in the Rosenthal & Henderson plot, Fig. 1D), and the estimated accuracy of angles was lower (0.236° in 1.66% glycerol vs. 0.368° in 20% glycerol). These data indicate that while glycerol does not necessarily prevent the ability to resolve a high-resolution reconstruction of a large, symmetric, and stable biological complex, it nonetheless negatively impacts the data quality.

**Figure 1.**
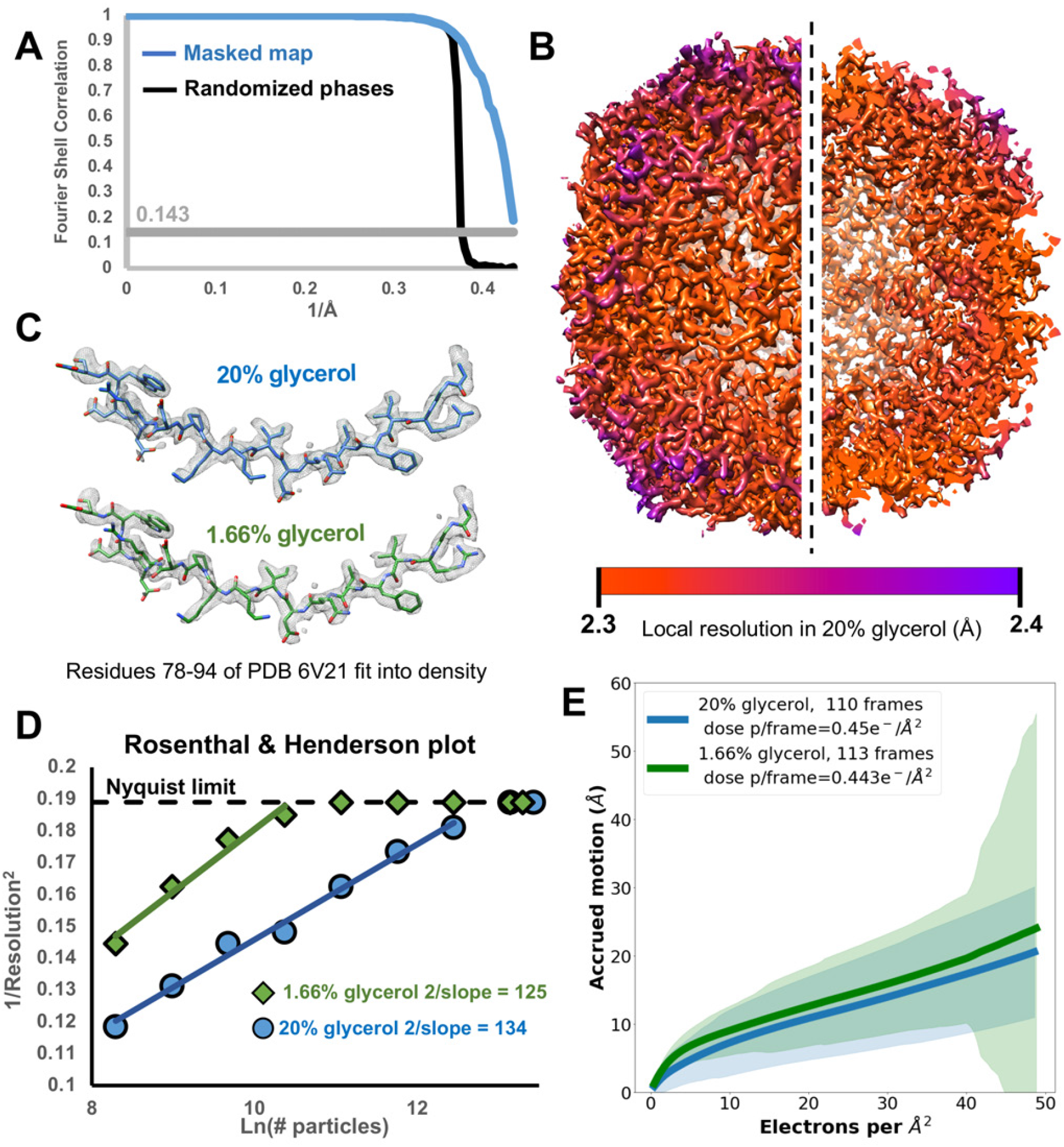
Apoferritin with 20% glycerol. **A**. Fourier shell correlation (FSC) between masked half maps (blue line) and between phase-randomized half maps (black line) from the apoferritin reconstruction in the presence of 20% glycerol. The grey line denotes the 0.143 correlation value. **B**. Reconstruction of apoferritin in the presence of 20% glycerol, colored by local resolution. The right half of the image is a cutaway showing the interior. **C**. Apoferritin model (PDB ID 6V21) fit into the EM density from reconstructions in 20% (blue) and 1.66% (green) glycerol. **D**. B-factor plot (Rosenthal & Henderson, 2003) showing the improvement of resolution as a function of the number of particles used for reconstruction in presence and absence of high glycerol concentration. **E**. Accrued beam-induced motion (per-frame summation) as a function of the number of electrons per Å^2^ received by the sample. Solid lines: average over all collected movies at a given dose. Colored shades: Standard deviation over all movies at a given dose.

### High concentrations of glycerol prevent the formation of optimally thin ice, lowering contrast

These two datasets offered an opportunity to perform a quantitative characterization of the impact that glycerol has on cryoEM data collection, and to identify the sources of data quality degradation. Notably, there was no indication of increased radiation damage in the form of bubbling present in any of the micrographs containing 20% glycerol, nor did the accrued motion significantly differ between the +/-glycerol samples (Fig. 1E). However, the images acquired in the presence of 20% glycerol showed qualitatively lower contrast than the samples with no added glycerol. We suspected that the likely source of this lowered contrast was the apoferritin particles being embedded in a thicker layer of ice when glycerol is present. The increased viscosity of the 20% glycerol sample was evident during the grid preparation process, as the blotting time required to obtain ice thin enough for data acquisition was increased nearly two-fold for the 20% glycerol sample (Figs. 2*a* and 2*b*). Longer blotting time is generally associated with thinner ice as the filter paper absorbs an increasing volume of sample solution. Despite this longer blot time, we were unable to attain sufficiently thin ice to produce a monolayer of apoferritin across the holes in the presence of 20% glycerol. This is evidenced by the presence of overlapping apoferritin particles in the micrographs and 2D averages (Fig. 2b and Supplementary Fig. 6).

**Figure 2.**
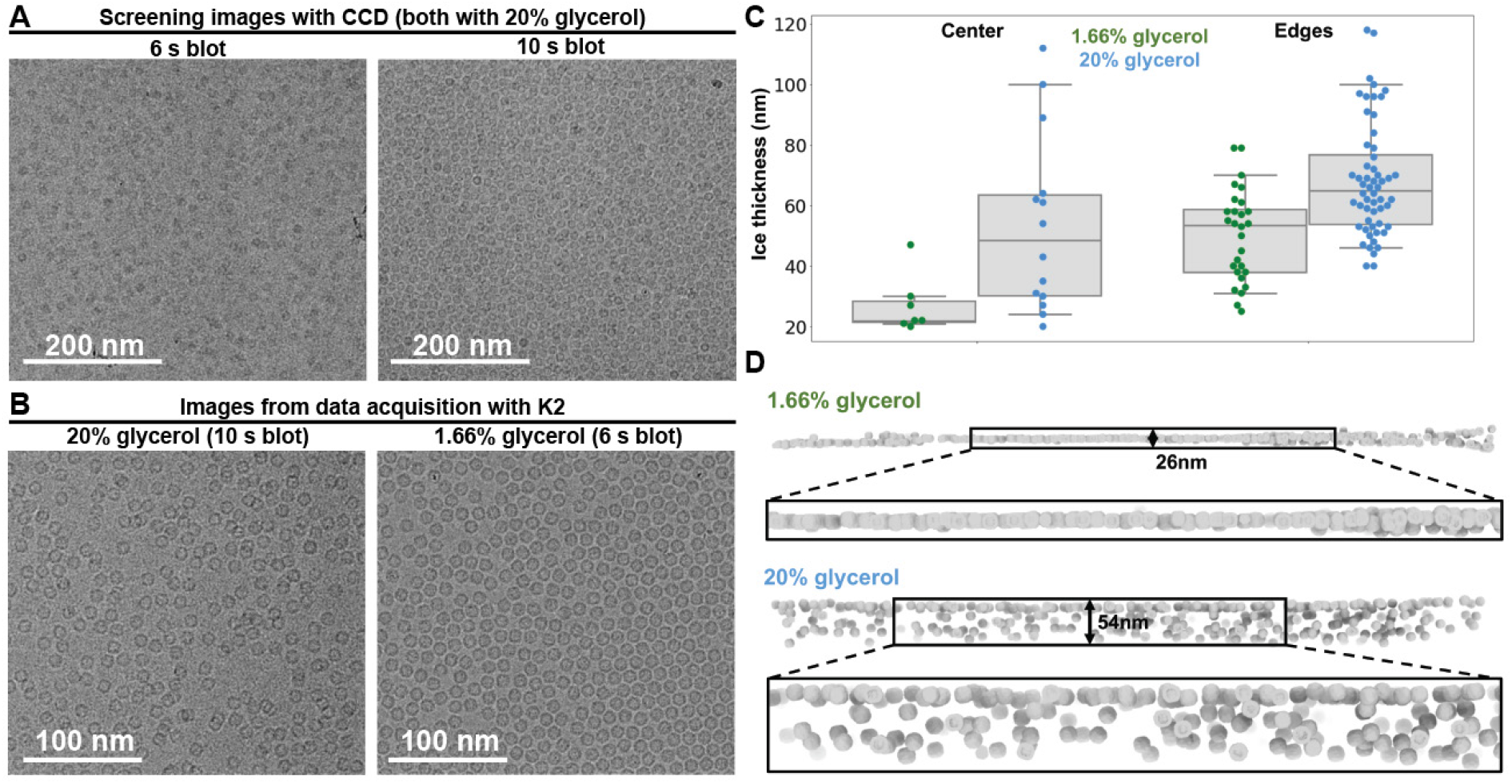
Assessing ice thickness of samples. **A**. Electron micrographs from sample screening, manually blotted for different times. **B**. Exemplar micrographs of samples with 1.66% and 20% glycerol. **C**. Ice thickness measurements obtained by electron tomography. Data points are shown as colored dots (horizontally stacked only to aid clarity) overlaid on a grey box that spans the two central quartiles and is crossed by a vertical line at the median. The whiskers span the range of data between the 5^th^ and 95^th^ percentiles. **D**. 3D renderings of ice cross sections from representative micrographs in + and – glycerol samples, generated by matching an apoferritin density template on the aligned tomograms and placing oriented density surface renderings as markers at high cross-correlation locations.

We further characterized the difference in ice thickness between the +/-glycerol samples of apoferritin using cryo electron tomography. Newly prepared apoferritin samples containing 1.66% and 20% glycerol were loaded into a Titan Krios operating at 300 keV and tomograms of the grid holes were acquired (7 tomograms of the 1.66% glycerol sample, 14 tomograms of the 20% glycerol sample). The tomogram analysis revealed ice in higher glycerol concentration samples was, on average, twice as thick. Notably, in the presence of 20% glycerol, the ice thickness at the center of the hole is more variable in different holes, while at 1.66% the ice at the center of the holes was reproducibly thin (Fig. 2c). Ideally, for single particle cryoEM analyses the ice thickness should only be slightly thicker than the diameter of the targeted particle in order to minimize the contribution of buffer molecules to the detected image. For this reason, images are routinely acquired at the center of the grid holes, where ice is generally thinnest and particles are often arranged as a single monolayer. In the presence of 1.66% glycerol, particles are mostly located in a single layer between the two surfaces of the air-buffer interface. In contrast, we identifed much fewer regions with comparably thin ice on the grid containing the 20% glycerol sample.

Intriguingly, the tomograms of the 20% glycerol grid confirmed apoferritin particles were located throughout the vitrified ice within the holes, as opposed to being adsorbed to the air-water interfaces, as observed previously in the absence of glycerol (Rice *et al*., 2018). This may be due to the increased viscosity of the buffer limiting the mobility of the particles during the blotting process, or perhaps by diminishing the hydrophobic interactions that promote adhesion of particles to the air-water interface. While this may benefit overall particle stability, this also results in the appearance of multiple overlapping layers of particles when viewed along the optical axis (Fig. 2*d* and Supplementary Figs. 6 and 9). Micrographs for single particle analysis were also acquired on the 20% glycerol sample that was used for tomographic data collection, and processing these data yielded a reconstruction with a reported resolution at Nyquist frequency (2.1 Å resolution) (Supplementary Fig. 8).

These data provide an explanation for the higher B-factors associated with the 20% glycerol sample. The increased ice thickness increases the likelihood of electron scattering due to interactions with the buffer molecules, detracting from the particle contrast in the images. The distribution of particles at different heights within the ice also results in the appearance of overlapping particles in the images, which can complicate accurate assignment of alignment parameters to each particle. Furthermore, the presence of particles at multiple heights in thick ice results in a distribution of defocus values corresponding to the particles, which decreases the accuracy of CTF estimation. Given the close proximity of particles to one another, it is also unlikely that per-particle CTF estimators will be able to assign defocus values to individual particles with high accuracy.

### Reconstruction of rabbit muscle aldolase at ∼3.3 Å in the presence of glycerol

Our apoferritin tests demonstrated that although glycerol increased ice thickness, thereby decreasing the overall quality of a single particle dataset, we were nonetheless able to determine a high-resolution structure of a stable, symmetric, 500 kDa protein complex. We next set out to test whether the addition of glycerol imposes resolution boundaries to smaller specimens, which are much more sensitive to ice thickness due to the lower overall electron scattering contribution of the biological specimen. To test the impact of 20% glycerol on a sample smaller than 200 kDa, we selected rabbit muscle aldolase, a protein that assembles into a 160 kDa homotetramer with D2 symmetry. Previously, we generated a reconstruction of this complex at ∼2.6 Å resolution using a 200 keV electron microscope, and to test the impact of glycerol we prepared the same sample with 20% glycerol using the same imaging parameters and microscope as used previously. Similarly to the apoferritin samples, a blotting time of 10 s was necessary to obtain sufficiently thin ice for data acquisition, which was ∼2 times longer than required for sample in the absence of glycerol (Wu *et al*., 2020). Despite a qualitatively noticeable decrease in contrast in our micrographs compared to those previously acquired without glycerol, we were able to obtain a ∼3.3 Å resolution in presence 20% glycerol (Figs 3*a*, 3*b* and Supplementary Fig. 10). Notably, the structural detail in our reconstruction was of sufficient quality to confidently assign side-chain identity in the core of the reconstruction (Fig. 3*c*). Analysis of the micrographs and frame alignment statistics showed, as for apoferritin, the added glycerol did not result in radiation-induced bubbling or any significant increases in beam-induced motion (Supplementary Fig. 11).

**Figure 3.**
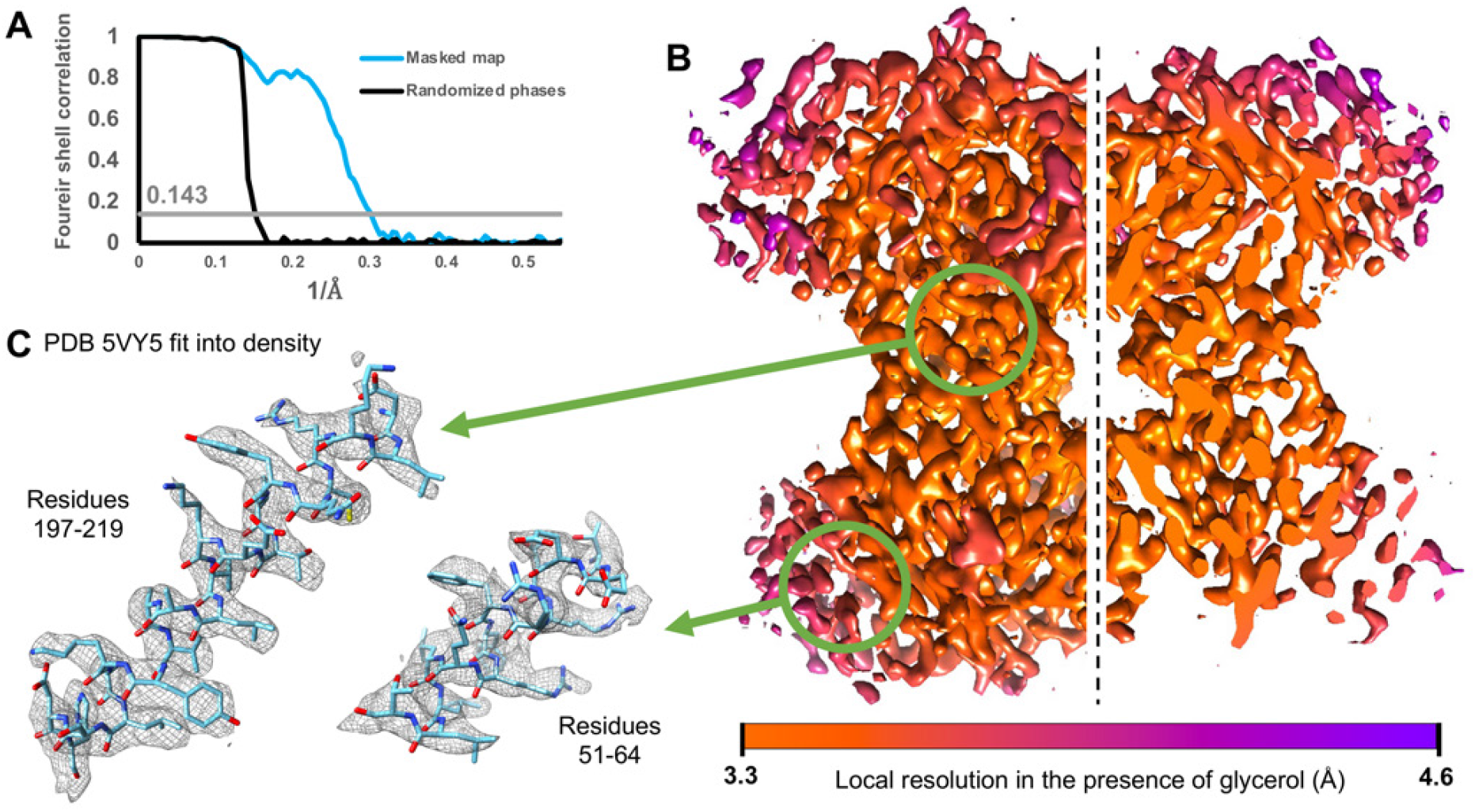
Aldolase in 20% glycerol. **A**. Fourier shell correlation between masked half maps (blue line) and between phase-randomized half maps (black line) from the aldolase density reconstruction in the presence of 20% glycerol. The grey line crosses the 0.143 gold-standard correlation value. **B**. Density reconstruction of aldolase from data acquired in the presence of 20% glycerol, colored by local resolution. **C**. Enlarged segments of aldolase density reconstruction from two areas with different local resolution. The PDB 5VY5 was aligned to the density and the corresponding atoms are shown inside the mesh.

We prepared additional aldolase grids with 20% glycerol to assess ice thickness by tomography on the 300 keV Krios, and confirmed that the ice was substantially thicker than that previously reported for aldolase prepared in the absence of glycerol (Rice *et al*., 2018), (Supplementary Fig. 12). Single particle data collected from these grids yielded a reconstruction at 4.6 Å resolution (estimated accuracy of angles: 6.82°, Supplementary Fig. 13). We do not believe the lower resolution of this reconstruction is due to imaging parameters or microscope, but due to the fact that the ice was likely thicker than for the 200 keV aldolase dataset, underscoring the challenge of consistently obtaining thin ice when glycerol is present at such high concentrations.

### Concluding remarks

Our results indicate that despite presenting challenges for optimal ice formation, introduction of glycerol molecules into the buffer does not abolish the ability to determine highresolution structures of biological specimens using single particle cryoEM. At the electron doses (50 and 80 e^-^ /Å^2^) and buffer compositions used in this study, we do not observe significant changes in beam-induced motion or radiation-induced bubbling for glycerol-containing samples compared to those with little or no glycerol. We do, however, confirm that the introduction of glycerol to the buffer lowers the overall quality of the data. We found that the lower contrast of glycerol-containing samples originates from the difficulty of generating thin ice that is comparable to samples lacking glycerol. Such issues with ice thickness can also arise from additives other than glycerol, such as DMSO, and ice thickness optimization is a necessary aspect of cryoEM sample prep. Sample volumes, ambient humidity, blotting times and forces could all be optimized to attain appropriate ice thickness. Of course, lower glycerol concentrations (e.g. 10% v/v) could potentially be used to attain thinner ice without substantially impacting macromolecular stability.

It is important to note that an often-overlooked benefit of thicker ice is the loosened confinement of large or dimensionally anisotropic particles. It is common for large particles to retreat to edges of holes where they are better accommodated by thicker ice, while particles in the centers of the holes are either denatured/broken, or forced to assume a preferred orientation that best accommodates the thinner ice. Thus, the limited thinning of ice offered by the addition of glycerol may aid in increasing the spatial and orientational distribution of particles in holes. Further, thicker ice may be preferable for fragile biological samples that denature at the air-water interface. Given that macromolecular targets requiring higher levels of glycerol for purification or functional assays likely fall under this category of fragile complexes, preservation in the thick ice promoted by 20% glycerol may be beneficial for structural stability and structure determination. Our tomograms show that the apoferritin particles are distributed throughout the vitrified buffer in holes, as opposed to being localized at the air-water interfaces, indicating that the addition of glycerol may protect a certain fraction of the sample from destructive hydrophobic interactions. Due to its small size, the same analysis was not possible for tomograms containing aldolase.

Lastly, we also note that direct plunging of an apoferritin sample containing 20% glycerol into liquid nitrogen seems to yield vitreous ice (Supplementary Fig. 14). As previously demonstrated by Taylor and Glaeser in the first studies of frozen hydrated catalase crystals (Taylor & Glaeser, 1974), the cryoprotective qualities of glycerol enable vitrification of the sample without the need for plunging into a cryogen with high heat capacity such as ethane or propane. This observation could conceivably pave the way to simpler workflows where manipulation of fragile grids and ice contamination are minimized (Tyler Engstrom *et al*., 2021).

We conclude glycerol should not be discarded as a cryoEM sample buffer additive, particularly for large, fragile complexes that are prone to disassembly or aggregation upon its removal. Further, the thicker ice and seemingly diminished interaction of particles with the air-water interface associated with the addition of glycerol may aid promoting a wider range of orientations in ice. We expect these findings to encourage new structural studies of samples that were previously not considered amenable for cryoEM analyses.

## Materials and Methods

### Sample preparation

Mouse apoferritin (heavy chain) was prepared as described previously (Wu *et al*., 2020). The protein was expressed in *E. coli* BL21(DE3)pLys, from a pET24a vector (Danev *et al*., 2019). Cells were grown to an OD_600_ of 0.5 at 37 °C in 500 mL of LB medium with shaking at 220 rpm. IPTG was then added to 1 mM to induce expression. Growth continued in the same conditions for 3 h, after which cells were harvested and pelleted at 4000 × *g* for 10 min at 4 °C. The pellet was resuspended in 20mL of lysis buffer (30 mM HEPES pH 7.5, 300 mM NaCl, 1 mM MgSO_4_) supplemented with 1 mg/ml lysozyme and cOmplete Protease Inhibitor Cocktail (Roche). Cells were lysed by sonication, and the resulting lysate clarified by centrifugation at 20000 × *g* for 30 min at 4 °C. The clarified lysate was heated for 10 min at 70 °C to precipitate endogenous *E. coli* proteins. Denatured proteins were pelleted by spinning at 20,000 × *g* for 15 min at 4 °C. Ammonium sulfate was added to the clarified lysate to a concentration of 60% (w/v), followed by gentle stirring on ice for 10 min. The precipitate was centrifuged at 14000 × *g* for 20 min. The resulting pellet was resuspended in 2 mL of cold PBS and dialyzed against Q1 buffer (30 mM HEPES pH 7.5, 1 mM DTT, 20 mM NaCl). After dialysis, the protein was further diluted by adding an equal volume of Q1 and loaded onto a HiTrap Q HP anion exchange chromatography column (GE Healthcare) equilibrated in buffer Q1. The column was washed with 4 volumes of Q1 and eluted with a 0 to 100% gradient of Q2 with SEC buffer (30 mM HEPES pH 7.5, 1 mM DTT, 500 mM NaCl) applied over 3 column volumes. Apoferritin eluted between 150 and 200 mM NaCl, which was confirmed by SDS-PAGE. The relevant fractions were pooled and concentrated to 10-20 mg/mL for loading on a Superdex 200 Increase 10/300 (GE Healthcare) size exclusion chromatography column equilibrated with 30 mM HEPES pH 7.5, 150 mM NaCl, 1 mM DTT. The peak fractions corresponding to apoferritin were pooled and concentrated to 15 mg/mL. For long-term storage, glycerol was added to 5% (v/v).

Rabbit muscle aldolase was prepared as described previously (Herzik *et al*., 2017). Lyophilized powder was purchased from Sigma-Aldrich and solubilized in 20 mM HEPES pH 7.5, 50 mM NaCl to a final concentration around 3 mg/ml. To remove aggregated protein the protein solution was filtered through a 0.2 µm filter and loaded on a Sepharose 6 10/300 (GE Healthcare) column equilibrated in solubilization buffer for size-exclusion chromatography. The fractions containing native, pure rabbit muscle aldolase were pooled and concentrated to 14.5 mg/mL.

### Cryo-electron microscopy sample handling and grid preparation

Prior to application on cryoEM grids, the apoferritin sample was diluted to 5 mg/mL using either SEC buffer (a final glycerol concentration of 1.66%, “no added glycerol” experiments), or SEC buffer supplemented with glycerol (“20% glycerol” experiments), such that the final glycerol concentration was 20% (v/v). 4 µL of apoferritin sample were applied on UltrAuFoil R1.2/1.3 300-mesh grids (Quantifoil Micro Tools GmbH) that had been freshly plasma cleaned for 7 seconds at 15 Watts (75% nitrogen/25% oxygen atmosphere) using a Solarus plasma cleaner (Gatan, Inc.). Grids were manually blotted using Whatman No. 1 filter paper and immediately plunge-frozen into liquid ethane cooled by liquid nitrogen using a custom-built manual plunger located in a cold room (≥95% relative humidity, 4 °C). For samples with 1.66% glycerol we blotted for 6 s. Expecting high glycerol concentrations to pose issues with ice thickness based on anecdotal evidence, we prepared grids of samples with 20% glycerol by blotting for 6, 8 and 10 s. After screening grids in cryogenic conditions (FEI F20 electron microscope operating at 200 keV, nominal magnification of 62,000x, pixel size 1.774 Å, using a Tietz 4kx4k CCD camera with a defocus of –1.6 µm using the Leginon automated electron microscopy package (Suloway *et al*., 2005), we decided to collect data on grids blotted for 6 s for the “no glycerol” condition and 10 s for the “20% glycerol” condition.

Prior to application on cryoEM grids, the aldolase stock was diluted to 1.6 mg/mL using buffer supplemented with glycerol, such that the final glycerol concentration was 20% (v/v). As with the apoferritin sample, 4 µL of aldolase sample were applied on UltrAuFoil R1.2/1.3 300-mesh grids (Quantifoil Micro Tools GmbH) that had been freshly plasma cleaned for 6 seconds at 15 Watts (75% nitrogen/25% oxygen atmosphere) using a Solarus plasma cleaner (Gatan, Inc.). Grids were manually blotted using Whatman No. 1 filter paper and immediately plunge-frozen into liquid ethane cooled by liquid nitrogen using a custom-built manual plunger located in a cold room (≥95% relative humidity, 4 °C). In order to optimize ice thickness in the presence of glycerol we prepared grids by blotting for 6, 8, 10 and 12 s. Following screening as done for the apoferritin samples, we decided to collect data on grids blotted for 10 s.

### CryoEM data acquisition for single particle analysis

For the grids imaged at 200 keV, movies of frozen-hydrated aldolase and apoferritin in the presence of 20% glycerol were collected using a Talos Arctica TEM (Thermo Fisher Scientific) with a field emission gun operating at 200 keV, equipped with a K2 Summit direct electron detector (Gatan, Inc.). Alignments were done as previously described to attain parallel illumination (Herzik *et al*., 2017, Lander *et al*., 2017), and image shift was used to image holes in a 4×4 array for apoferritin, and a 2×2 array for aldolase, using stage movement only to shift between subsequent arrays. Movies were collected in counting mode (1.15 Å/pixel and 0.91 Å/pixel, respectively) at a dose of 50 e^-^ /Å^2^ and 80 e^-^ /Å^2^, respectively (110 frames of 100 ms, and 96 frames of 200 ms, respectively) and a nominal defocus of –1.0 µm and –1.3 µm, respectively. Movies of apoferritin in the absence of glycerol were collected in the same imaging conditions as apoferritin in the presence of 20% glycerol, with the only difference being the collection of 113 frames of 100 ms.

For the grids imaged at 300 keV, movies of frozen-hydrated apoferritin and aldolase in the presence of 20% glycerol were collected using a Titan Krios TEM (Thermo Fisher) with a field emission gun operating at 300 keV, equipped with a K2 Summit direct electron detector (Gatan, Inc). Image shift was used to image holes in a 4×4 array for apoferritin, and a center-of-five array for aldolase, using stage movement only to shift between subsequent arrays. Movies were collected in counting mode (1.026 Å/pixel and 1.045 Å/pixel, respectively) at a dose of 50 e^-^ /Å^2^ (65 frames of 100 ms, and 50 frames of 200 ms, respectively) and defocus ranges between of -0.5 and -1.5 µm for apoferritin, and between -1.25 and -1.75 µm for aldolase. Tomography data was acquired on a small region of these same samples prior to data collection for single particle analysis (see section 6 for details).

All cryoEM data for single particle analysis were acquired using the Leginon automated data collection software (Suloway *et al*., 2005) and pre-processed in real-time using the Appion package (Lander *et al*., 2009). Pre-processing consisted of beam-induced motion correction and contrast transfer function (CTF) estimation. Frame alignment and dose weighting were done using MotionCor2 (Zheng *et al*., 2017) without binning, on 5×5 patches, 7 iterations, a global B factor of 500 and a local B factor of 100. CTF estimation was performed using CTFFIND4 (Rohou & Grigorieff, 2015) on the aligned but non-dose-weighed frames, using the following parameters: 1024 field size, 0.07 amplitude contrast, 50 Å minimum resolution, 4 Å maximum resolution. Real time CTF estimation was only used for monitoring data quality during collection.

### Processing apoferritin in low glycerol at 200 kV

We used RELION 3.1 (Zivanov *et al*., 2018) for data analysis of apoferritin without added glycerol (Supplementary Fig. 1). We aligned dose-weighted movie frames using MotionCor2 from within RELION 3.1 in order to enable later Bayesian polishing, and estimated initial CTF parameters using CTFFIND4 from within RELION 3.1. For particle picking, we began by manually picking ∼50 particles from 19 randomly selected micrographs. We extracted the picked particles in boxes of 192 pixels, without binning (1.15 Å/pixel), and classified them into five 2D classes. We selected the one class that most clearly resembled apoferritin as a template for template-based picking in the AutoPick RELION 3.1 node. This resulted in 876,503 picks from 1,707 micrographs. 2D classification (5 classes, Tau=2, 25 iterations) of these picks produced 3 classes with clear secondary structure features, which contained 875,694 particles. EMD-21024 was low-pass filtered to 30 Å to serve as the initial model for auto-refinement (O symmetry imposed) of these particles, which yielded a reconstruction at 3.9 Å resolution using an FSC cut-off of 0.143. Particles were grouped by image shift followed by per-micrograph defocus and astigmatism refinement, combined with beam tilt refinement, yielding a reconstruction at 2.4 Å. Bayesian polishing, followed by 3D classification without alignment (O symmetry imposed, 4 classes, Tau=20, 100 iterations) and further CTF refinement increased resolution to 2.3 Å (FSC = 0.143).

### Processing apoferritin in 20% glycerol at 200 kV

We used RELION 3.1 for data analysis of apoferritin in the presence of 20% glycerol (Supplementary Fig. 2). We aligned dose-weighted movie frames using MotionCor2 from within RELION 3.1 in order to enable later Bayesian polishing, and estimated initial CTF parameters using CTFFIND4 from within RELION 3.1. We initially manually picked 2,261 particles from 20 randomly selected micrographs. We extracted the picked particles in boxes of 192 pixels, without binning (1.15 Å/pixel), and classified them into five 2D classes. We used the four classes that most clearly resembled apoferritin as templates for template-based picking in the AutoPick RELION 3.1 node. This resulted in 830,042 picks from 2,254 micrographs. 2D classification (5 classes, Tau=2, 25 iterations) of these picks produced two classes with clear secondary structure features, which contained 829,923 particles. EMD-21024 was low-pass filtered to 15 Å and used as an initial model to auto-refine (O symmetry imposed) these particles to yield a reconstruction at 4.2 Å resolution according to FSC at 0.143. After grouping by image shift, CTF refinement (per-particle defocus, per-micrograph astigmatism, and beam tilt) and 3D refinement (O symmetry imposed) the resolution improved to 2.9 Å at FSC = 0.143. A round of 3D classification (O symmetry imposed, 6 classes, Tau=4, 30 iterations) and subsequent class selection by visual inspection yielded a smaller particle set (683,154 particles) which, after 3D refinement (O symmetry imposed), CTF refinement, Bayesian polishing and further CTF refinement, yielded the final reconstruction at 2.3 Å (FSC=0.143).

### Processing apoferritin in 20% glycerol at 300 Kv

We used RELION 3.1 for data analysis of apoferritin in the presence of 20% glycerol (Supplementary Fig. 3). We aligned dose-weighted movie frames using MotionCor2 from within RELION 3.1 and estimated initial CTF parameters using CTFFIND4 from within RELION 3.1. We manually picked 277 particles from 5 randomly selected micrographs, and extracted the particles binned by 4 (4.12 Å/pixel, 48 pixel box size), and ran 2D classification requesting a single class. This class was used as a template for template picking, resulting in ∼1.8 million picks that were extracted binned by 4 (4.12 Å/pixel, 48 pixel box size) and subjected to 2D classification (50 classes, Tau=2, 25 iterations). Selection of 19 high quality 2D classes, in which secondary structure was visible, yielded a particle stack of 778,762 particles from 1,859 micrographs. An initial model generated in Cryosparc v2.14.2 (Punjani *et al*., 2017) during previous exploratory work was used as a reference model for 3D auto-refinement (O symmetry imposed) of the particle stack. As 3D classification failed to generate distinguishable classes, the whole particle stack was re-centered and re-extracted at full resolution (1.03 Å/pixel, 192 pixel box size). Refinement (O symmetry imposed) of the unbinned stack resulted in a reconstruction at a nominal resolution of 2.9 Å at FSC = 0.143. The particles were grouped by image shift and iteratively CTF refined with beam tilt correction and per particle defocus refinement. Subsequent auto-refinement (O symmetry imposed) improved the nominal resolution to 2.1 Å at FSC = 0.143.

### Processing aldolase in 20% glycerol at 200 kV

We used RELION 3.1 and Appion (Lander *et al*., 2009) for data analysis of aldolase in the presence of 20% glycerol (Supplementary Fig. 4). We aligned dose-weighted movie frames using MotionCor2 from within RELION 3.1 and then imported them to CryoSPARC for 2D analyses. Subsequent 3D classification and refinement was done in RELION 3.1. We initially selected particles using DoG Picker 2 (Voss *et al*., 2009) on the initial 76 micrographs after they were aligned and dose-weighted in real-time using the Appion launcher. These particles were extracted and used as input to Appion’s Iterative multivariate statistical analysis/multi-reference projection routine to create 2D class averages for template-based particle picking using FindEM (Roseman, 2004), on all 741 micrographs. The resulting 323,571 particle coordinates were imported to RELION 3.1 and extracted binned by 4 (3.64 Å/pixel) in a final box of 64 pixels. We performed four successive runs of 2D classification (50 classes, Tau=1, E-step resolution limited to 8Å, 30 iterations), at each iteration discarding only class averages that clearly did not contain particles. We re-centered and extracted the remaining 322,648 particles binned by 2 (1.82 Å/pixel) in a 128 pixel box and performed one more round of 2D classification where we only kept classes where secondary structure elements were visible and radial streaks from noise were minimal. We re-centered and re-extracted (no binning, 256 pixel box) this 142,300-particle set, subjected it to 3D auto-refinement (D2 symmetry imposed) starting from EMD-8743 low-pass filtered to 15Å, and obtained a reconstruction of aldolase with a reported resolution of 4.8 according to FSC = 0.143. Grouping by image shift and subsequent runs of CTF refinement, first refinement of beam tilt only, and subsequently of defocus and astigmatism, followed by auto-refinement yielded a reconstruction at 4.2 Å resolution (FSC = 0.143). We performed 3D classification without alignment (4 classes, Tau=8, 25 iterations), and selected particles from the highest resolution class. 3D auto-refinement using these 28,588 particles, followed by CTF refinement (defocus, astigmatism, and beam tilt) improved resolution after auto-refinement (D2 symmetry imposed) to 4.0 Å. Bayesian polishing, followed by CTF refinement (defocus, astigmatism, beam tilt, and trefoil) yielded a final reconstruction after auto-refinement (D2 symmetry imposed) at 3.3 Å resolution (FSC = 0.143).

### Processing aldolase in 20% glycerol at 200 kV

We used RELION 3.1 and CryoSPARCv3.1.0 (Punjani *et al*., 2017) for data analysis of aldolase in the presence of 20% glycerol (Supplementary Fig. 5). We aligned dose-weighted movie frames using MotionCor2 from within RELION 3.1 and then imported them to CryoSPARC for 2D classification, subsequent 3D classification and refinement were done in RELION 3.1. We used the pyem (Asarnow *et al*., 2019) library to convert CryoSPARC particle data output files to RELION 3.1 star files. We imported the aligned and dose-weighted micrographs in CryoSPARC, ran patch CTF estimation and selected micrographs containing CTF estimates reporting signal at 5 Å resolution or better (1870 of 2591 total micrographs). We then performed template picking using low-pass filtered projections of EMD-8743 as templates, yielding 1,255,598 particles that were extracted and input to CryoSPARC 2D classification. We performed two rounds of 2D classification. After the first run of 2D classification we selected all particles from classes that remotely resembled aldolase, even if the classes did not contain high-resolution features. Particles comprising these classes were used for a subsequent 2D classification run, after which only particles contained in classes bearing discernible secondary structure elements were selected. The coordinates of these 143,234 particles were re-centered and re-extracted without binning using a 256 pixel box size using RELION 3.1. The initial refinement from these particles yielded a reconstruction at 6.5 Å resolution (FSC = 0.143). We then removed duplicates, re-centered and re-extracted the particles, after which 3D auto-refinement yielded a reconstruction at 6.0 Å resolution. We then performed 3D classification without alignment (6 classes with Tau=6), from which we isolated a single class where secondary structure elements were discernible and radial streaking was minimal. This class contained 21,155 particles. Refinement using this particle set yielded a 5.6 Å resolution reconstruction. Grouping by image shift followed by beam tilt refinement and 3D auto-refinement improved the resolution to 5.3 Å, and a final refinement round using “Mask individual particles with zeros = No” yielded a reconstruction at 4.5 Å resolution (FSC = 0.143).

### Accrued beam-induced motion calculations

To calculate the accrued beam-induced motion we used the global shift data collected in the star files produced by RELION 3.1 after running MotionCor2 from within the RELION 3.1 interface. Only the micrographs containing particles from the final refinement set were used for this analysis (1,858 out of 2,261 for the apoferritin in 20% (v/v) glycerol dataset collected at 200 keV, 1,434 out of 1,707 for the apoferritin in 1.66% (v/v) glycerol dataset collected at 200 keV, and 735 out of 741 for the aldolase in 20% (v/v) glycerol dataset collected at 200 keV). To obtain the accrued motion per frame, we calculated the distance between the location in the current frame and the previous frame and summed it to the total previous traveled distance. The dose at each data point is calculated by the dose per frame multiplied by the frame number.

### Cryo-electron tomography data acquisition

Tilt series for apoferritin and aldolase samples were collected using a Thermo Fisher Titan Krios TEM at 300 keV, equipped with a Gatan K2 Summit direct electron detector. Tomography data was acquired on a small region of the grid, which was later avoided for single particle data collection. Data acquisition was done using SerialEM (Mastronarde, 2003). Tilt series were acquired using a sequential tilting scheme, starting at 0° and increasing to +60° at 2° increments, then returning to 0° and increasing to -60° at 2° increments. Each tilt series was collected with a nominal defocus of -6 to -8 µm for apoferritin and -10µm for aldolase samples. Each tilt was acquired as movies in counting mode using a per-frame dose of 0.9 e^-^ /Å^2^, and a pixel size of 3.73 Å. For aldolase, we acquired tilt series on two sets of 6 holes each, arranged on two transects on neighboring grid squares where particles were visible on imaging conditions used for single particle data acquisition. For apoferritin, we acquired 7 tilt series on a sample without added glycerol and 14 tilt series on a sample with glycerol in the buffer, on squares that were not used for single particle data collection but of similar quality.

### Cryo-electron tomography data analysis

Image stacks from apoferritin and aldolase were binned by a factor of 4 prior to tomogram alignment using the newstack application from IMOD (Kremer *et al*., 1996). Tomogram reconstruction was done in the eTomo (Mastronarde, 1997) IMOD module, following the pipeline from coarse alignment to fine and final alignments. Ice thickness measurements from reconstructed tomograms was performed as previously described (Rice *et al*., 2018). In brief, the binned tomograms were oriented in “3dmod slicer view” in IMOD for optimal viewing of the X-Z axis, with the top and bottom particle layers (i.e. the air-water interface) roughly parallel to the field of view. These particle layers on the air-water interface were used as indicators of the ice boundary. Occasionally ice contamination deposited on the ice surface further confirmed the location of the ice boundary. The 3dmod IMOD module was used to measure ice thickness at the hole edges and in the center, as delineated by the distance between the top and bottom particle layers.

The particle space-filling models embedded in ice shown in Fig. 2 and Supplementary Fig. 9 were created using dynamo (Castaño-Díez *et al*., 2012). The final density obtained by single particle analysis of the same sample was used as template for template matching in the aligned tomograms using dynamo. In order to prepare the density to be used as a template, we inverted the contrast using EMAN2’s (Tang *et al*., 2007) e2proc3d.py script, and down-sampled to the same pixel size as the tomograms using RELION 3.1 (Zivanov *et al*., 2018) auxiliary application relion image handler. A surface rendering of the same density, generated with dynamo, was then placed at each cross-correlation peak in the tomogram volume only in locations where cross-correlation values were above 0.33. To verify the depicted particles were not located at correlation maxima derived from noise we plotted the raw tomogram data of 100 randomly selected particles and visually confirmed they resembled apoferritin.

### Direct plunging in liquid nitrogen of an apoferritin sample in presence of 20% glycerol

The grids were plasma cleaned and protein samples were prepared as described above. Blotting was done from the side, using Whatman No. 1 filter paper, as normally done for negative stain. Plunging was done by hand, simply submerging grids completely in liquid nitrogen. These frozen hydrated samples were imaged at liquid nitrogen temperature on a Tietz 4kx4k CCD camera with a defocus of –2.5 µm using the Leginon automated electron microscopy package, in a FEI F20 electron microscope operating at 200 keV, at a nominal magnification of 62,000x (pixel size 1.774 Å).

### CryoEM map rendering and local resolution coloring

CryoEM maps were rendered using UCSF Chimera (Pettersen *et al*., 2004). Local resolution calculations were done in RELION 3.1, and the outputs used for local resolution coloring in Chimera.

## Data availability

All reconstructions were deposited to the Electron Microscopy Data Bank (EMDB) and are accessible with the following IDs: Apoferritin in 1.6% glycerol acquired at 200 keV: EMD-24795; Apoferritin in 20% glycerol acquired at 200 keV: EMD-24796; Apoferritin in 20% glycerol acquired at 300 keV: EMD-24797; Aldolase in 20% glycerol at 200 keV: EMD-24798; Aldolase in 20% glycerol at 300 keV: EMD-24799. Datasets are being uploaded to the Electron Microscopy Public Image Archive (EMPIAR)

## Acknowledgements

The authors thank Dr. Mengyu Wu for providing the motion correction data for the aldolase sample without glycerol and useful discussion, as well as Dr. Benjamin Barad for his help with tomography data acquisition. We thank Dr. Anette Schneemann for input on the manuscript. B.B. is supported by a Postdoctoral Fellowship from the George E. Hewitt Foundation for Medical Research. M.H. is supported by the Dennis and Marsha Dammerman Fellowship of the Damon Runyon Cancer Research Foundation. G.C.L. is supported by the National Institutes of Health (NIH) R21AG067594 and R01AG061697.

## Supplementary Materials

**Supplementary Figure 1.**
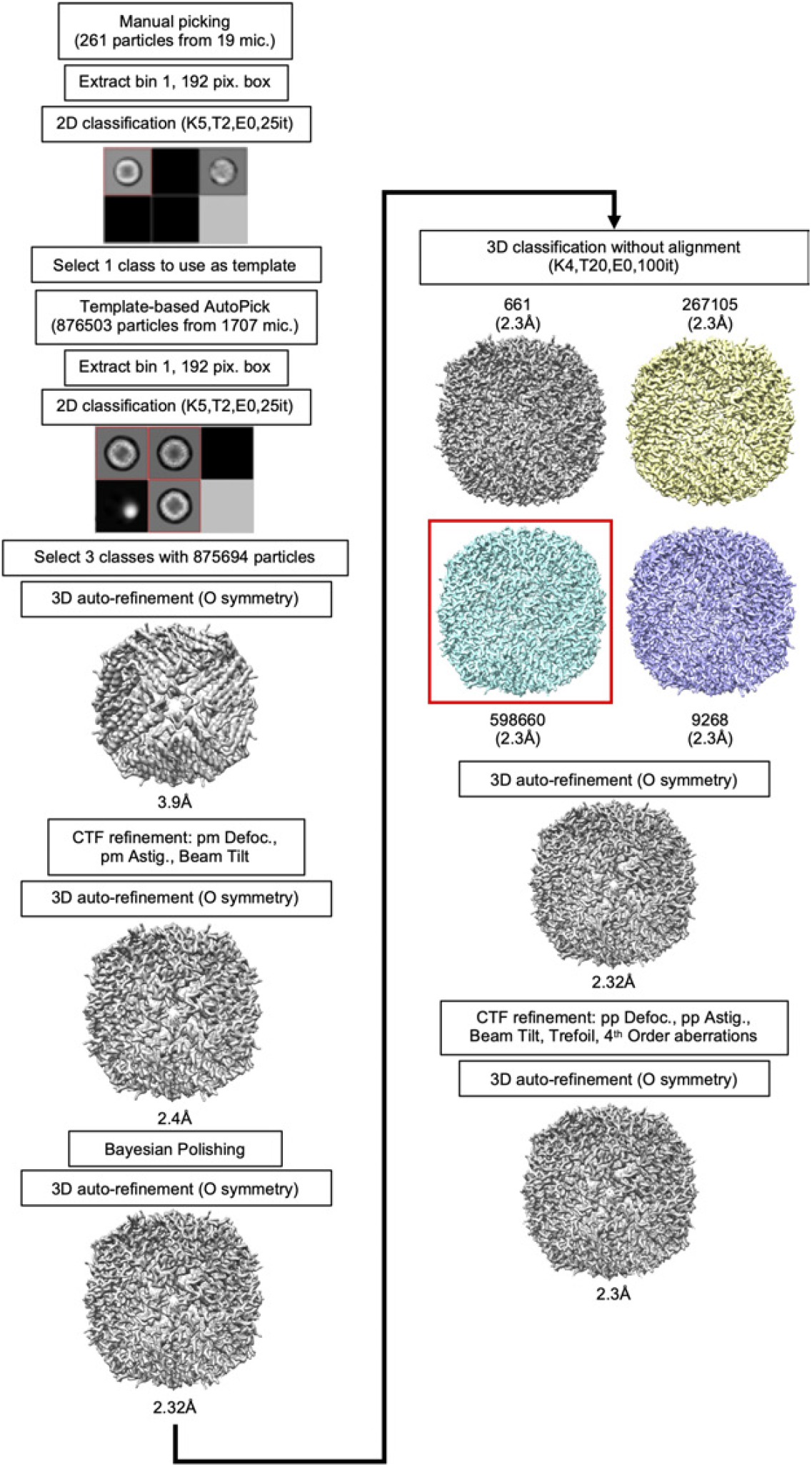
Data analysis workflow for apoferritin in 1.66% glycerol buffer imaged at 200KeV. In 2D and 3D classification steps, the selected classes are marked with a red box. Empty classes in 2D jobs are represented by dark squares. Relevant parameters for each RELION node are listed in each box.

**Supplementary Figure 2.**
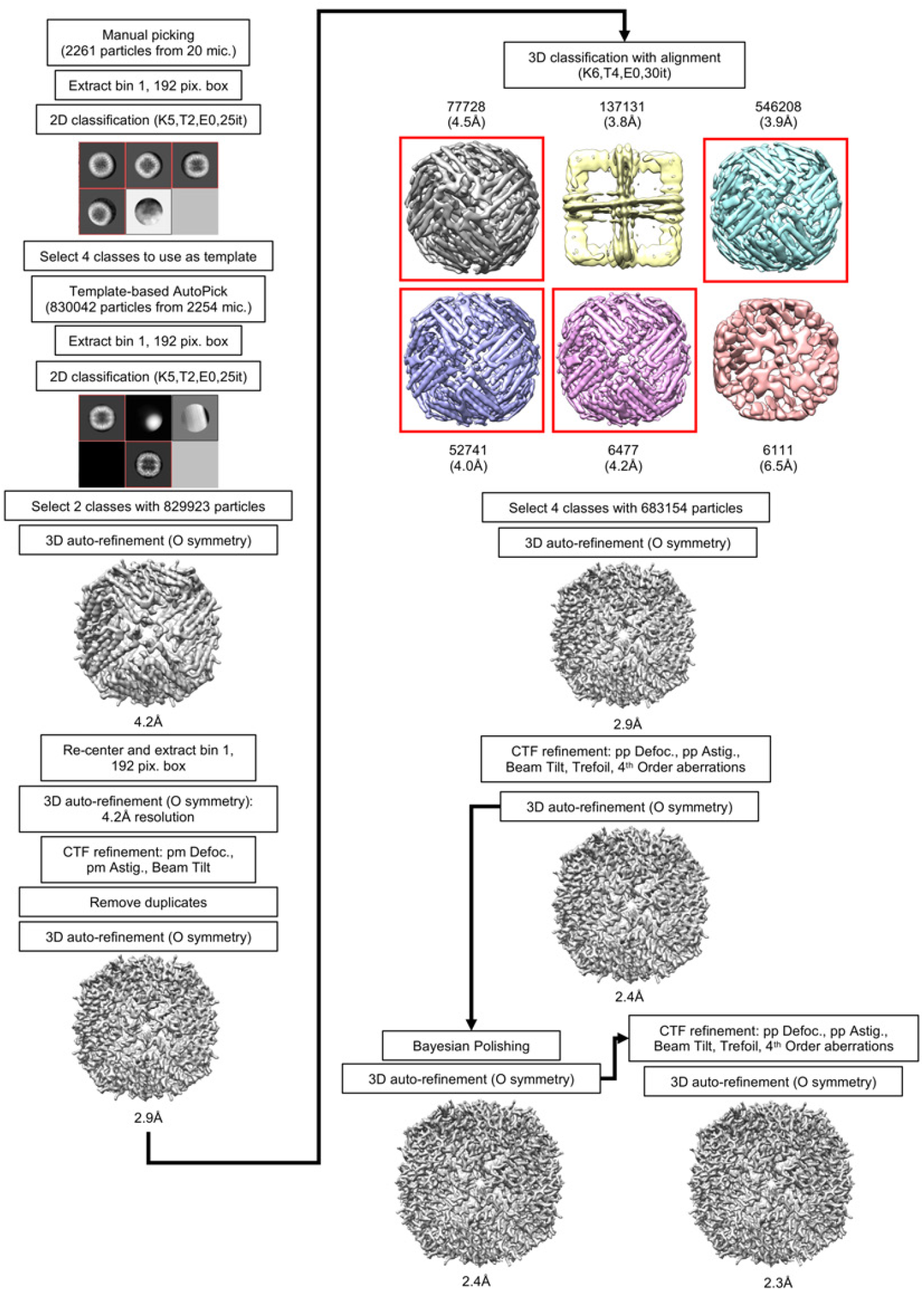
Data analysis workflow for apoferritin in 20% glycerol buffer imaged at 200KeV. In 2D and 3D classification steps, the selected classes are marked with a red box. Empty classes in 2D jobs are represented by dark squares. Relevant parameters for each RELION node are listed in each box.

**Supplementary Figure 3.**
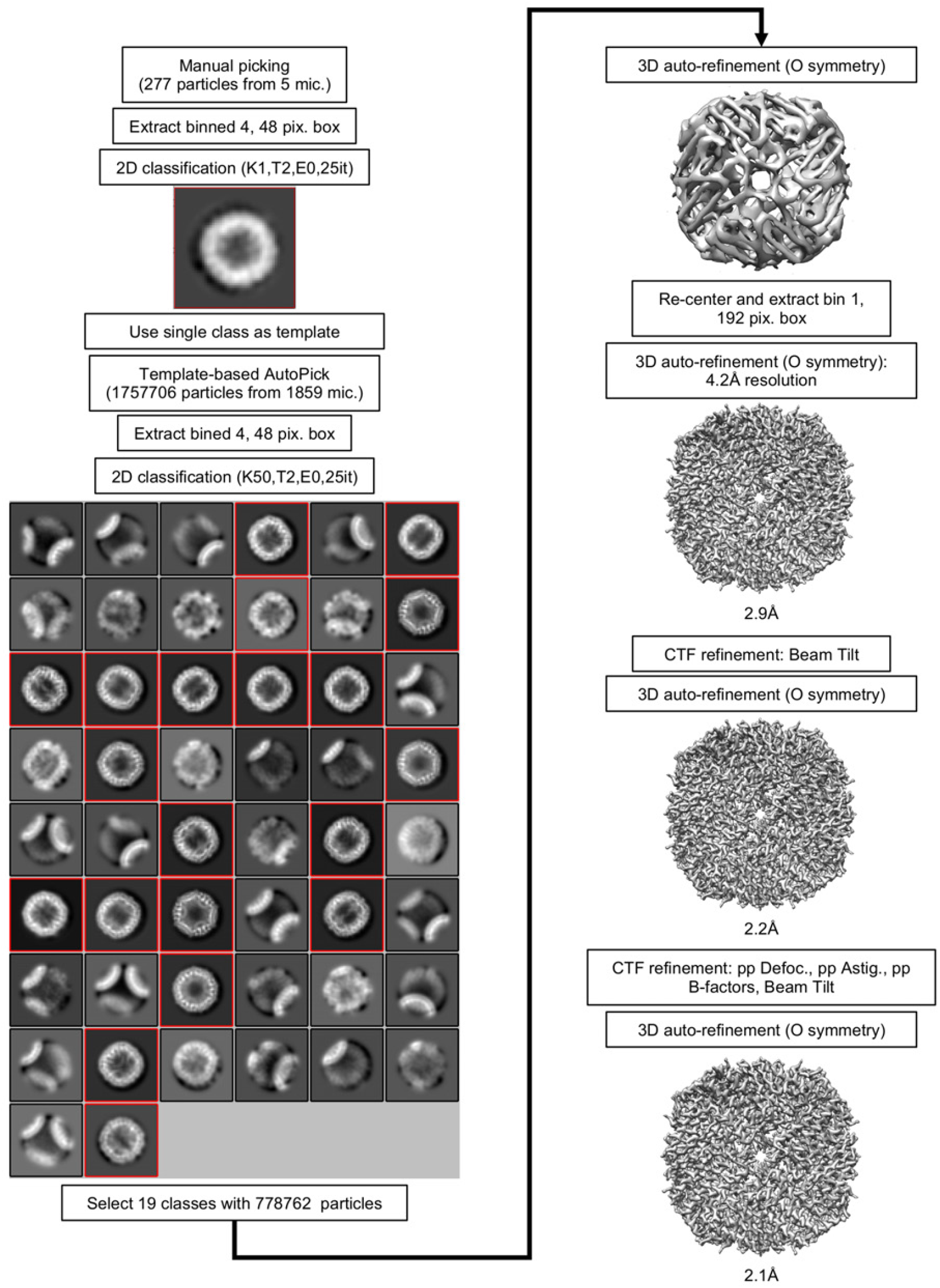
Data analysis workflow for apoferritin in 20% glycerol buffer imaged at 300KeV. In the 2D classification step, the selected classes are marked with a red box. Relevant parameters for each RELION node are listed in each box.

**Supplementary Figure 4.**
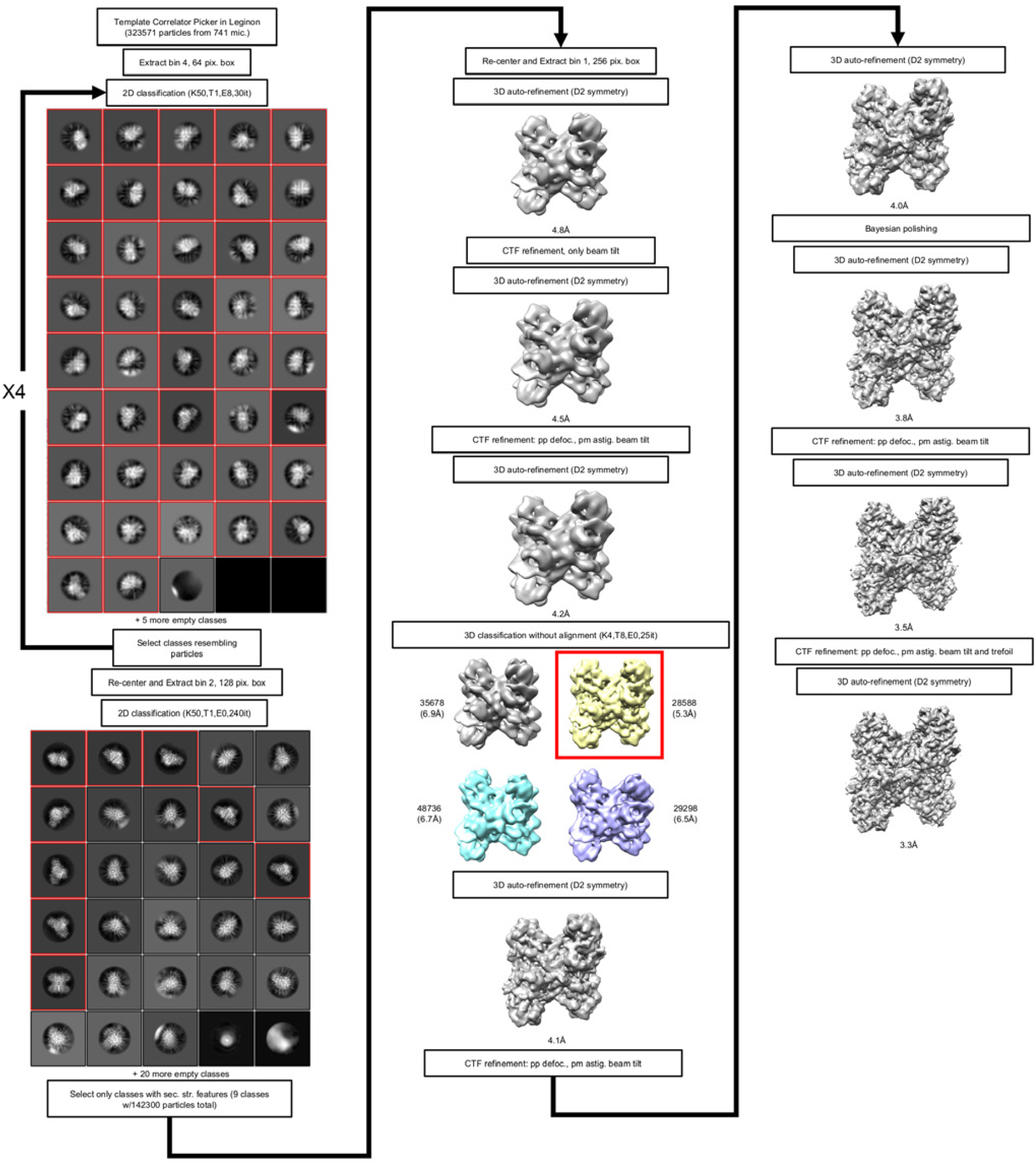
Data analysis workflow for aldolase in 20% glycerol buffer imaged at 200KeV. In 2D and 3D classification steps, the selected classes are marked with a red box. Empty classes in 2D jobs are represented by dark squares. Relevant parameters for each RELION node are listed in each box.

**Supplementary Figure 5.**
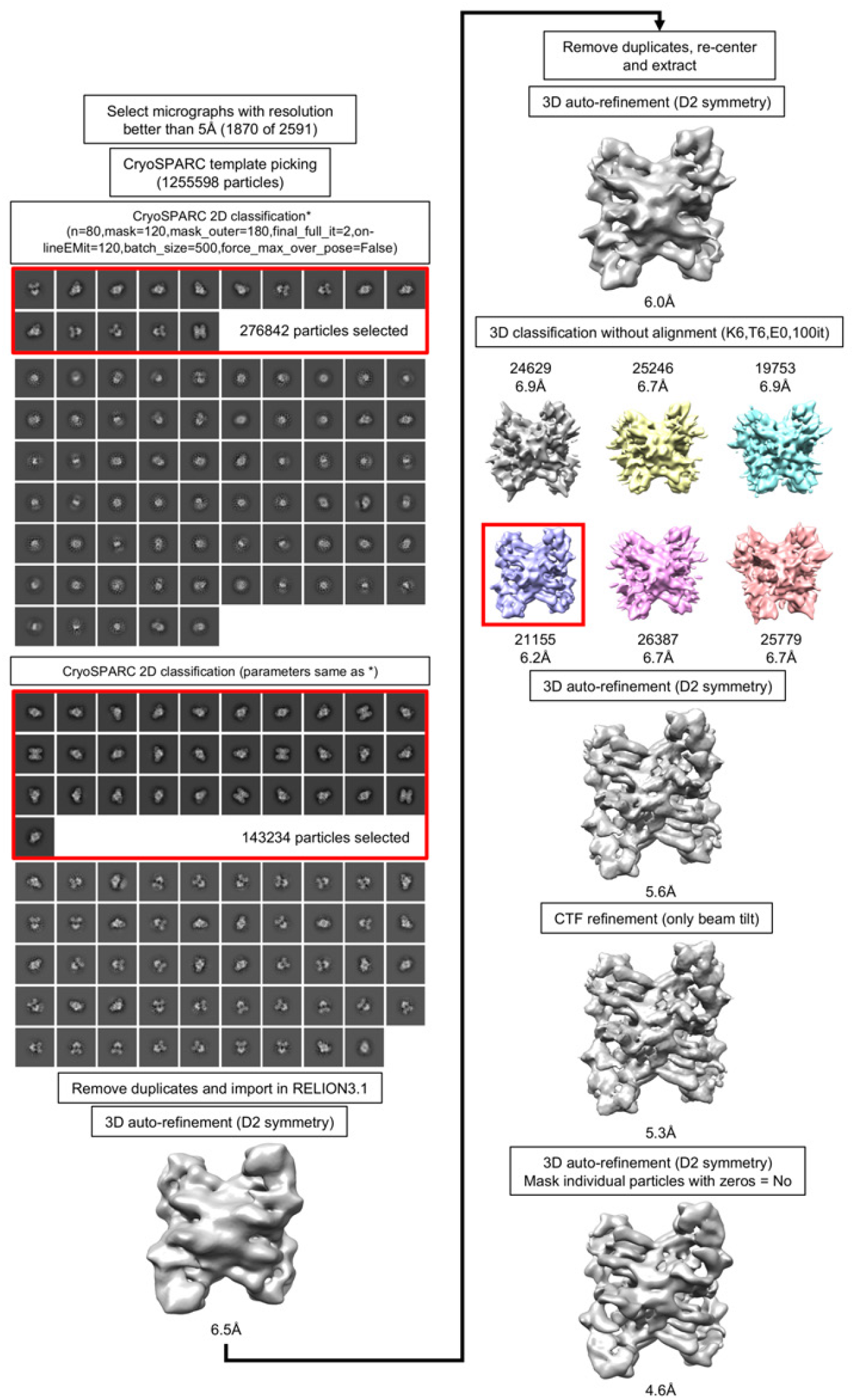
Data analysis workflow for aldolase in 20% glycerol buffer imaged at 300KeV. In 2D and 3D classification steps, the selected classes are marked with a red box. Empty classes in 2D jobs are represented by dark squares. Relevant parameters for each RELION node are listed in each box.

**Supplementary Figure 6.**
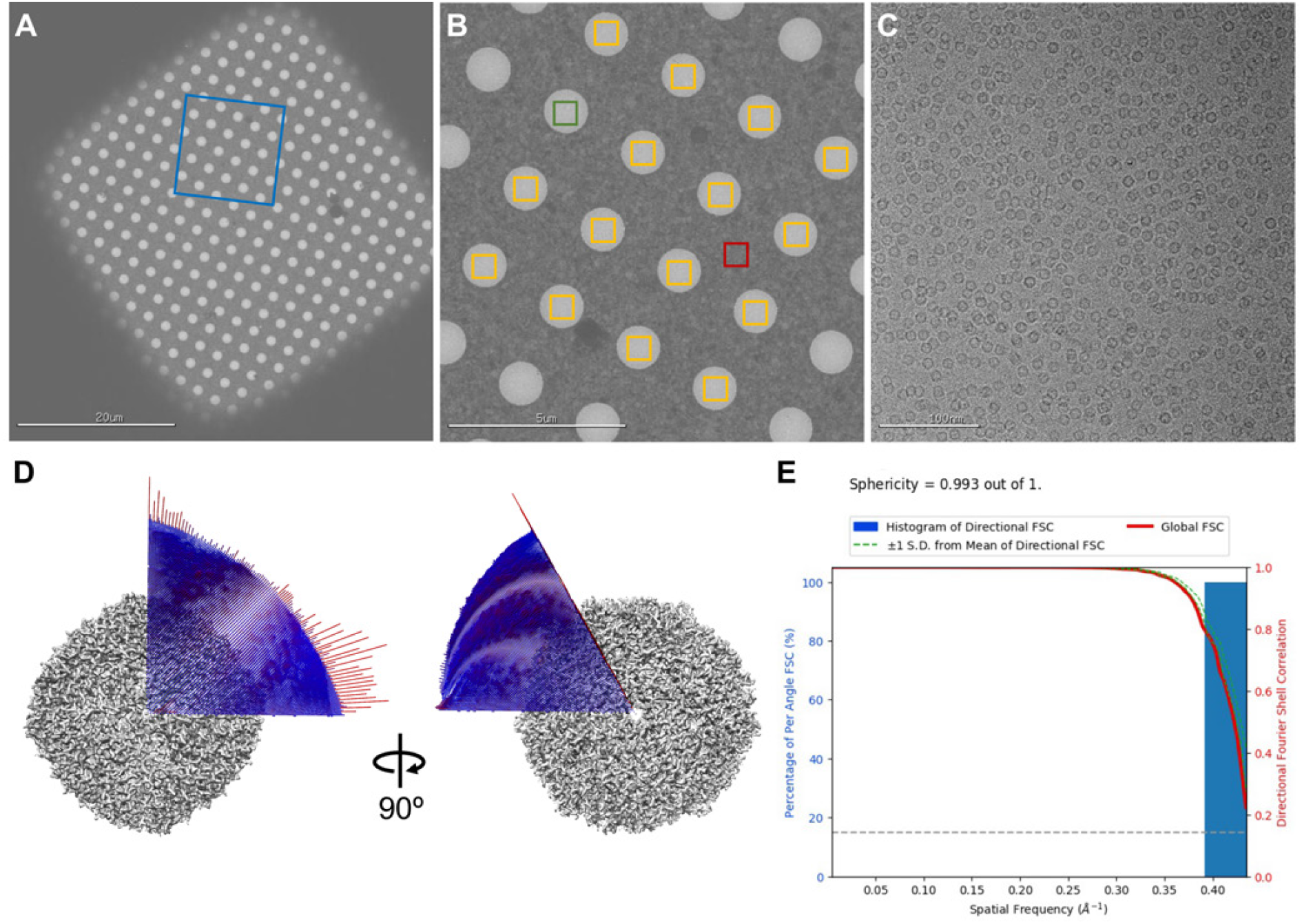
Exemplar micrographs and additional final reconstruction metrics for apoferritin imaged at 200KeV in 20% glycerol buffer. **A**. Exemplar grid square, a blue frame marks where micrograph (B) was acquired. **B**. An exemplar “hole” magnification micrograph, with a red square showing where real focus estimation was done, and yellow squares showing where high-magnification micrographs were acquired for single particle analysis. A green square shows where micrograph (C) was taken. **C**. An exemplar micrograph used for single particle picking and analysis. **D**. Angular distribution of views for the final reconstruction done assuming O symmetry. **E**. Directional FSC curves showing how resolution varies by orientation (Tan *et al*., 2017).

**Supplementary Figure 7.**
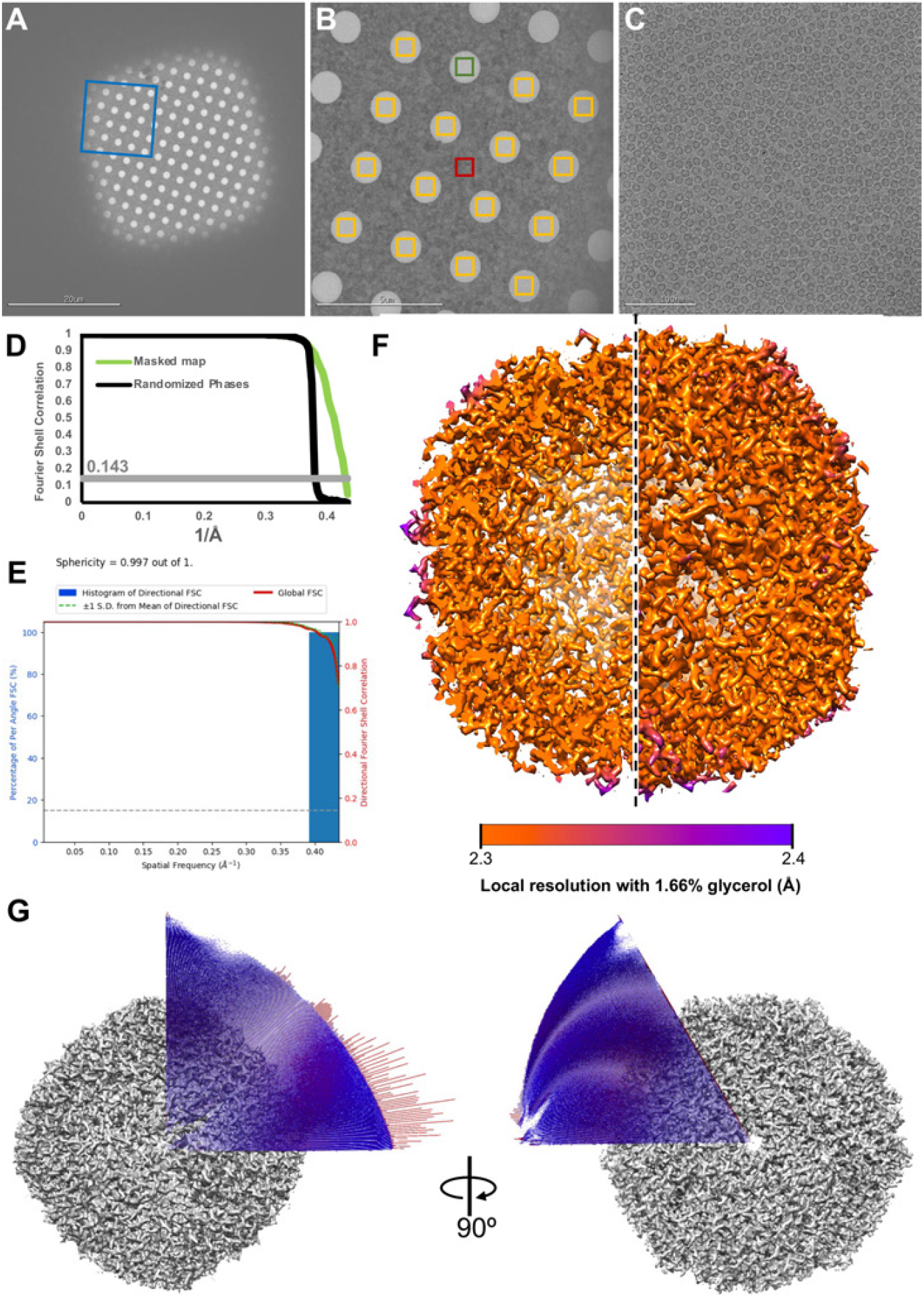
Exemplar micrographs and additional final reconstruction metrics for apoferritin imaged at 200KeV with 1.66% glycerol. **A**. Exemplar grid square, a blue frame marks where micrograph (B) was acquired. **B**. An exemplar “hole” magnification micrograph, with a red square showing where real focus estimation was done, and yellow squares showing where high-magnification micrographs were acquired for single particle analysis. A green square shows where micrograph (C) was taken. **C**. An exemplar micrograph used for single particle picking and analysis. **D**. Fourier shell correlation curve for the masked reconstruction and for randomized phases. **E**. Directional FSC curves showing how resolution varies by orientation (Tan *et al*., 2017). **F**. Reconstruction colored by local resolution, calculated by RELION 3.1 Local Resolution node (Zivanov *et al*., 2018). **G**. Angular distribution of views for the final reconstruction done assuming O symmetry.

**Supplementary Figure 8.**
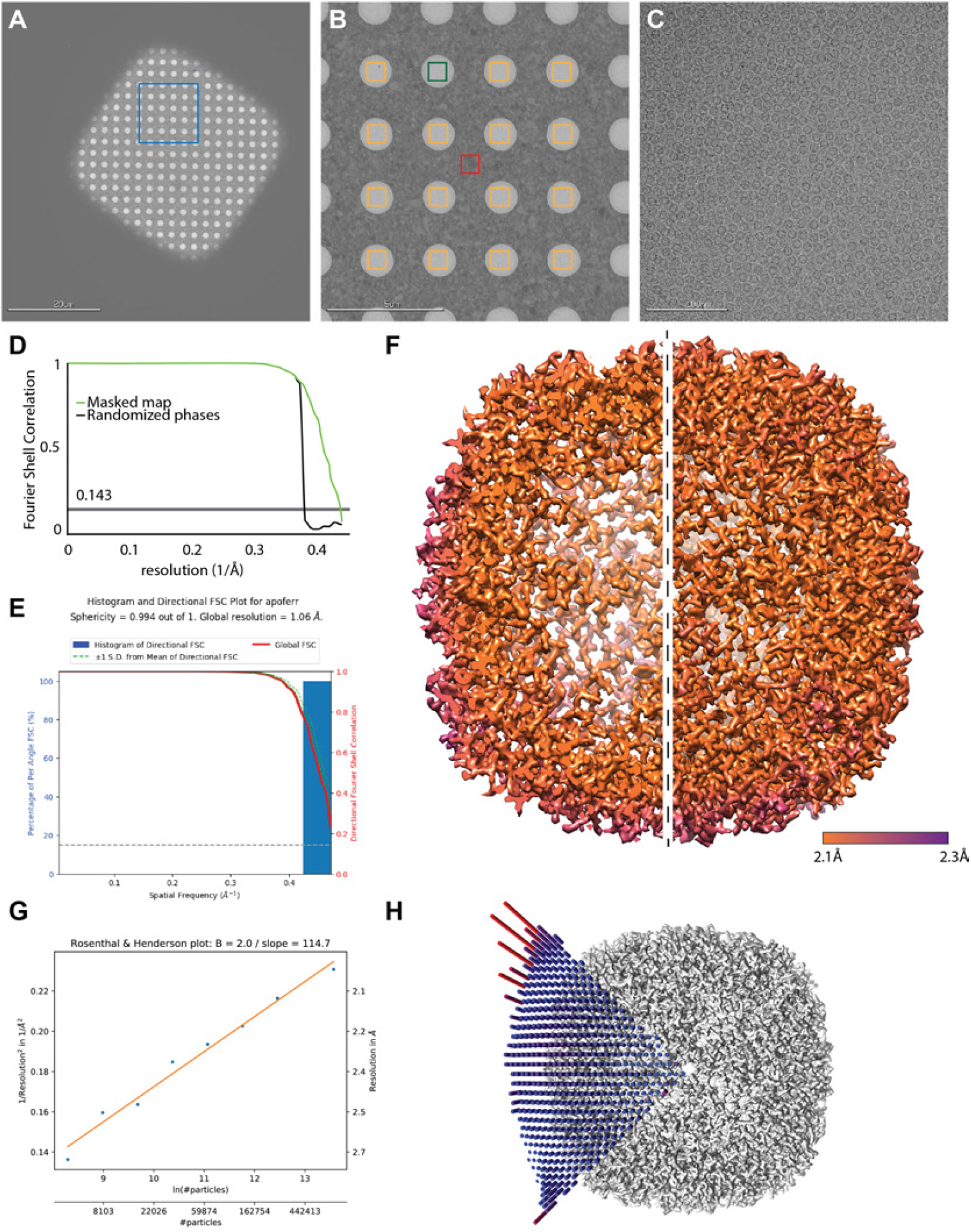
Exemplar micrographs and additional final reconstruction metrics for apoferritin imaged at 300KeV in 20% glycerol buffer. **A**. Exemplar grid square, a blue frame marks where micrograph (B) was acquired. **B**. An exemplar “hole” magnification micrograph, with a red square showing where real focus estimation was done, and yellow squares showing where high-magnification micrographs were acquired for single particle analysis. A green square shows where micrograph (C) was taken. **C**. An exemplar micrograph used for single particle picking and analysis. **D**. Fourier shell correlation curve for the masked reconstruction and for randomized phases. **E**. Directional FSC curves showing how resolution varies by orientation (Tan *et al*., 2017). **F**. Reconstruction colored by local resolution, calculated by RELION 3.1 Local Resolution node (Zivanov *et al*., 2018). **G**. B-factor plot showing how reconstruction resolution improves as larger subsets of the final particle set are used for 3D auto-refinement (Rosenthal & Henderson, 2003), generated using the bfactor plot.py script provided together with RELION 3.1 source code. **H**. Angular distribution of views for the final reconstruction done assuming O symmetry.

**Supplementary Figure 9A.**
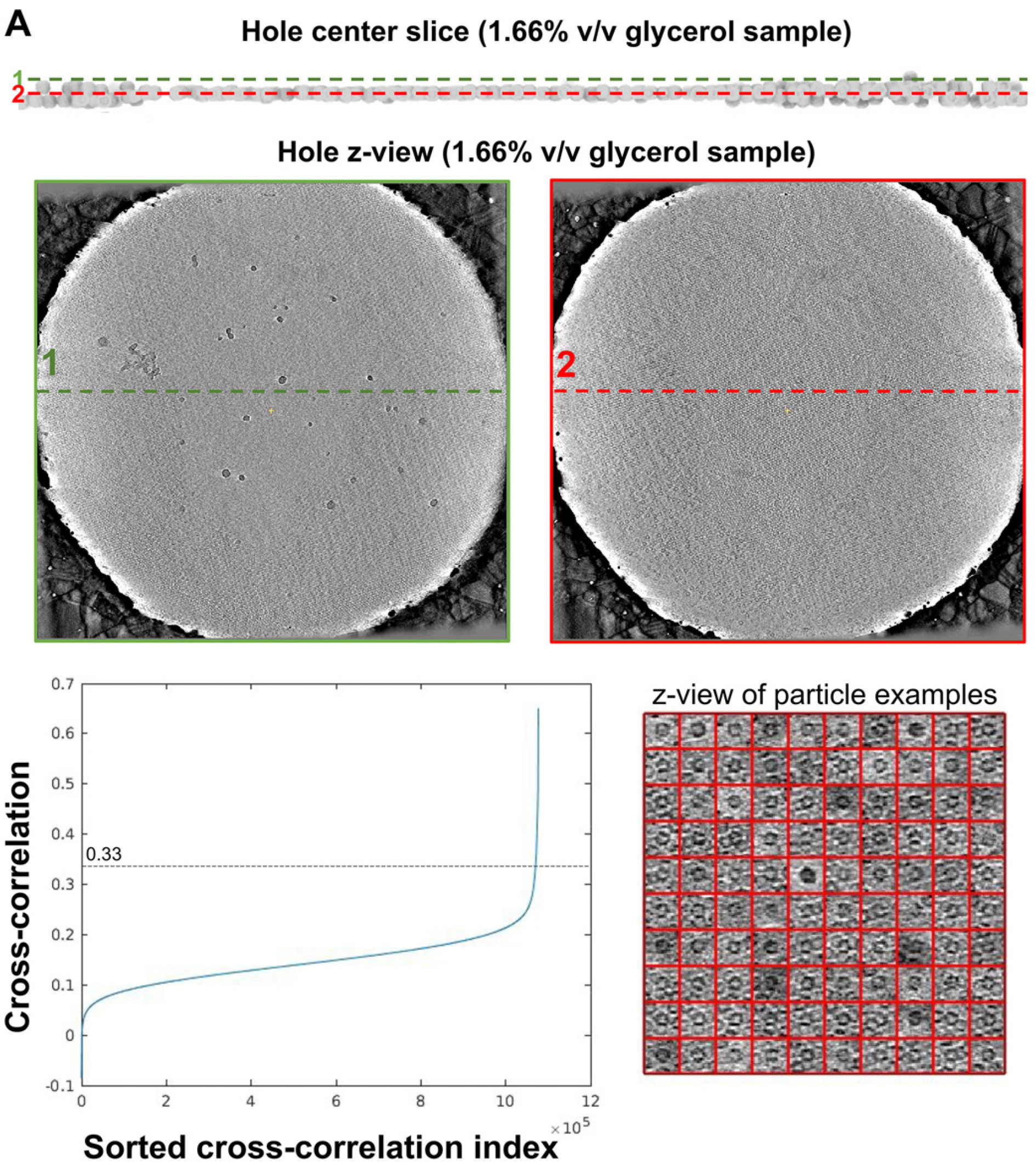
Distribution of particle layers in apoferritin samples with low glycerol concentration. An exemplar hole from the 1.66% glycerol sample, showing two parallel planes in the aligned tomogram at different ice depths (center) labeled on a schematic profile (top) derived from template matching using an apoferritin model. The bottom left panel shows cross-correlation values as a function of peak index sorted by cross-correlation, the labeled line at 0.33 cross-correlation indicates the threshold above which correlation peaks were used to construct the particle distribution model on the top. The bottom right panel shows 100 randomly-picked particles from the set used to construct the particle distribution model on the top, indicating the overwhelming majority of picks are clearly apoferritin particles.

**Supplementary Figure 9B.**
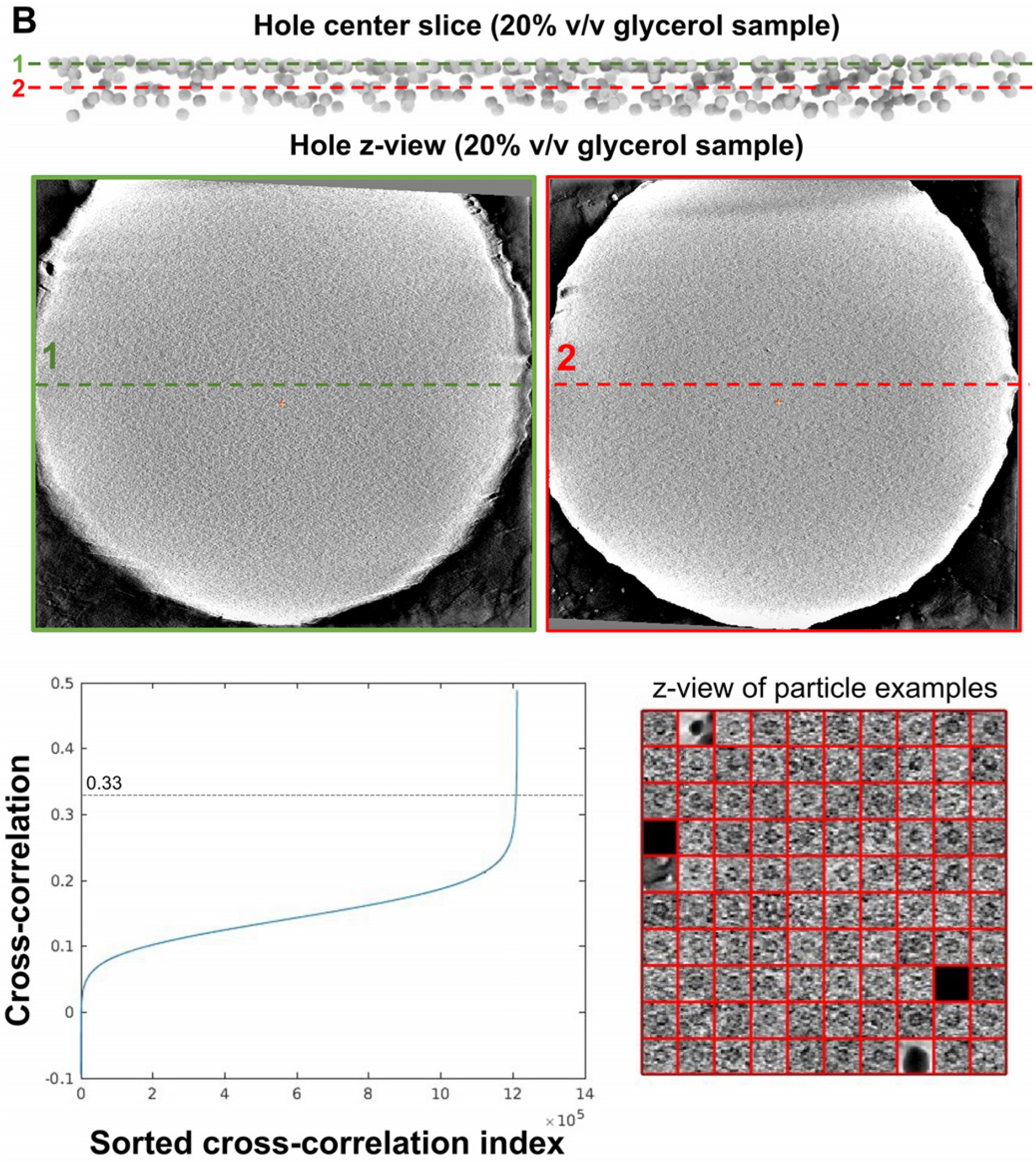
Distribution of particle layers in apoferritin samples with high glycerol concentration. An exemplar hole from the 20% glycerol sample, showing two parallel planes in the aligned tomogram at different ice depths (center) labeled on a schematic profile (top) derived from template matching using an apoferritin model. The bottom left panel shows cross-correlation values as a function of peak index sorted by cross-correlation, the labeled line at 0.33 cross-correlation indicates the threshold above which correlation peaks were used to construct the particle distribution model on the top. The bottom right panel shows 100 randomly-picked particles from the set used to construct the particle distribution model on the top, indicating the overwhelming majority of picks are clearly apoferritin particles.

**Supplementary Figure 10.**
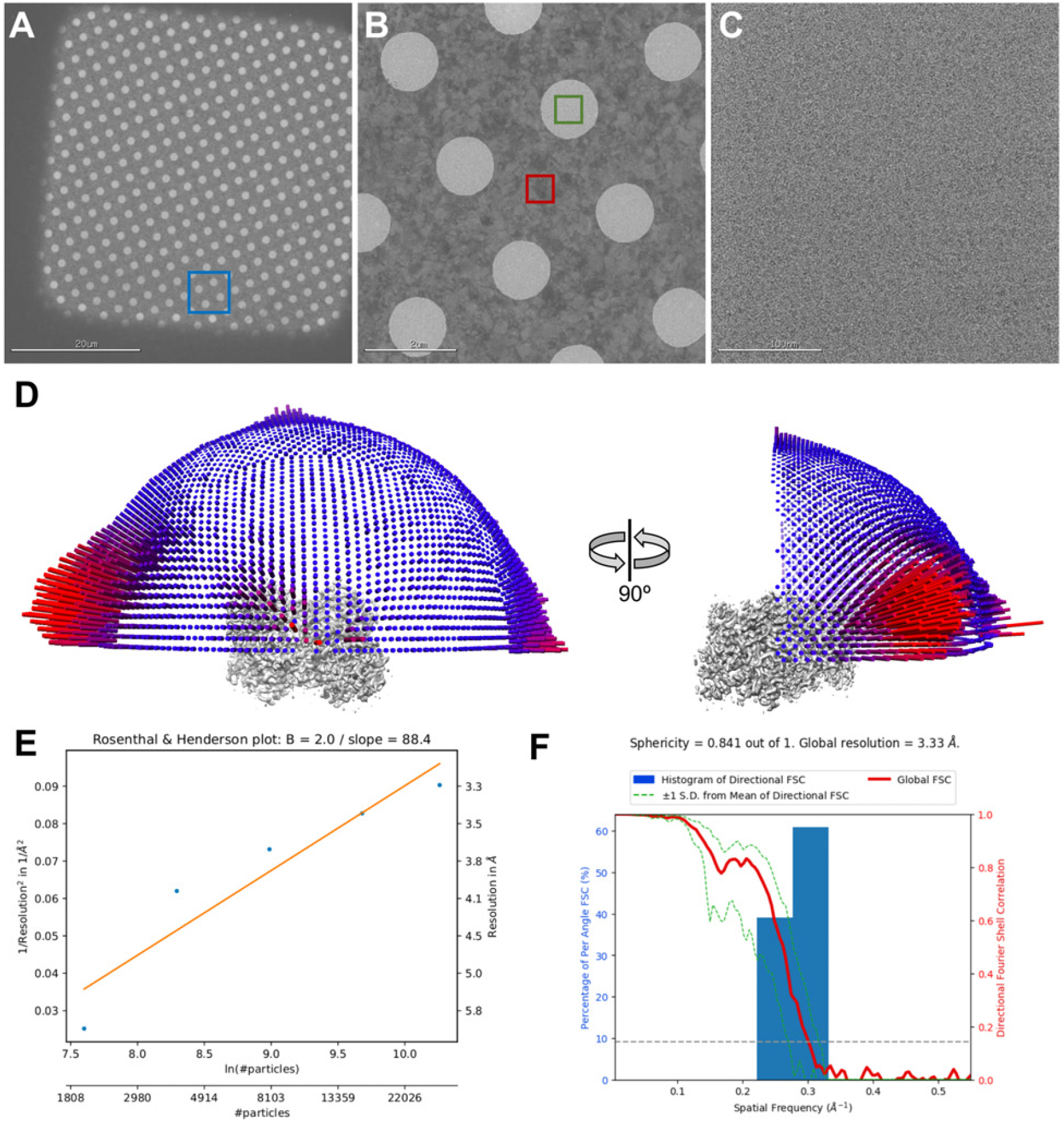
Exemplar micrographs and additional final reconstruction metrics for aldolase imaged at 200KeV in 20% glycerol buffer. **A**. Exemplar grid square, a blue frame marks where micrograph (B) was acquired. **B**. An exemplar “hole” magnification micrograph, with a red square showing where real focus estimation was done, and yellow squares showing where high-magnification micrographs were acquired for single particle analysis. A green square shows where micrograph (C) was taken. **C**. An exemplar micrograph used for single particle picking and analysis. **D**. Fourier shell correlation curve for the masked reconstruction and for randomized phases. **E**. Directional FSC curves showing how resolution varies by orientation (Tan *et al*., 2017). **F**. Reconstruction colored by local resolution, calculated by RELION 3.1 Local Resolution node (Zivanov *et al*., 2018). **G**. B-factor plot showing how reconstruction resolution improves as larger subsets of the final particle set are used for 3D auto-refinement (Rosenthal & Henderson, 2003), generated using the bfactor plot.py script provided together with RELION 3.1 source code. **H**. Angular distribution of views for the final reconstruction done assuming D2 symmetry.

**Supplementary Figure 11.**
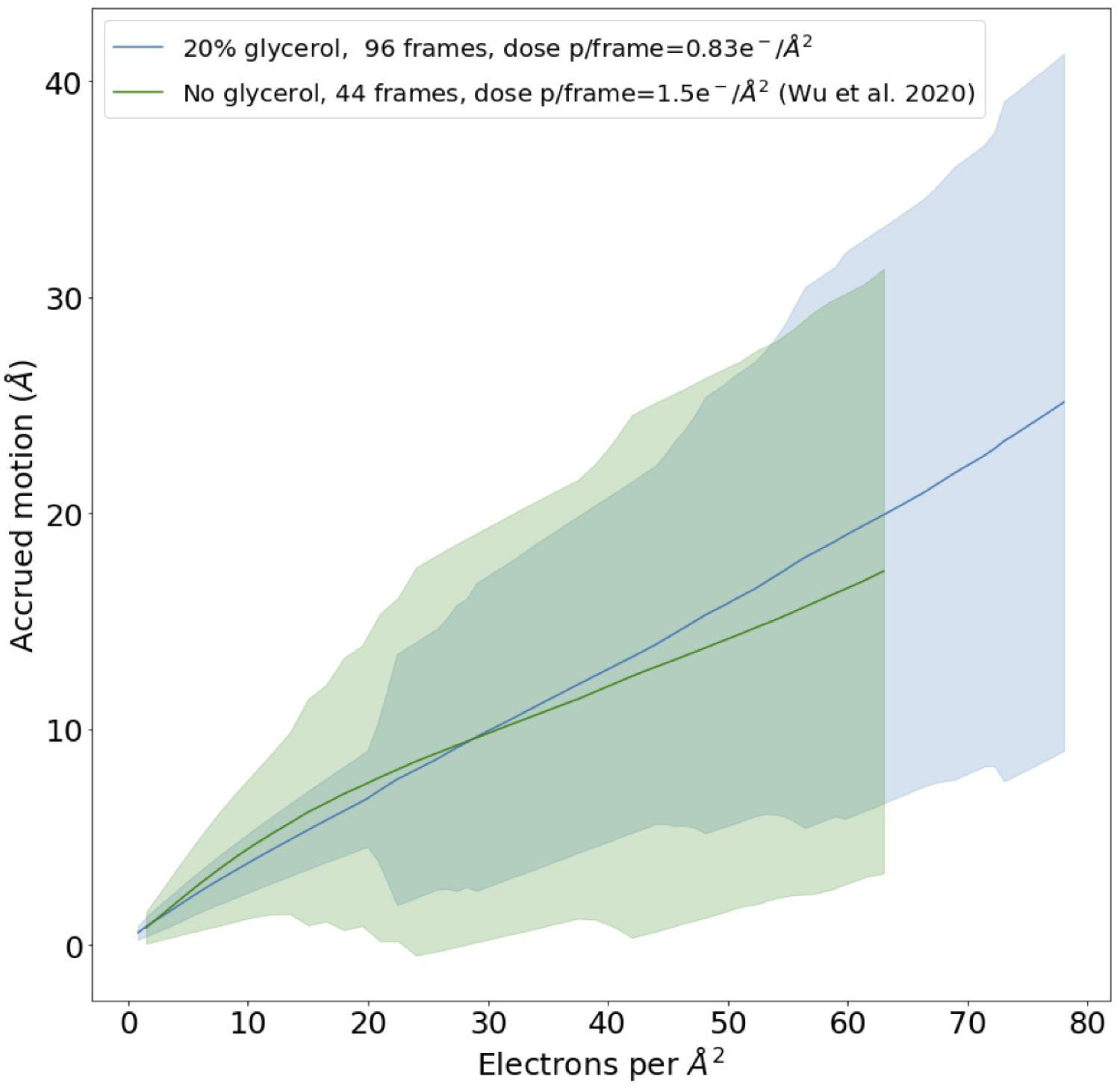
Accrued beam-induced motion for aldolase imaged at 200KeV in 20% glycerol buffer compared to a previous publication. Accrued beam-induced motion (per-frame summation) as a function of the number of electrons per Å^2^ received by the sample. Our aldolase sample in 20% glycerol buffer, imaged at 200KeV (blue), compared to a previously published aldolase dataset without glycerol, with a similar imaging setup at 200KeV – green (Wu *et al*., 2020). Solid lines: average over all collected movies at a given dose. Colored shades: Standard deviation over all movies at a given dose.

**Supplementary Figure 12.**
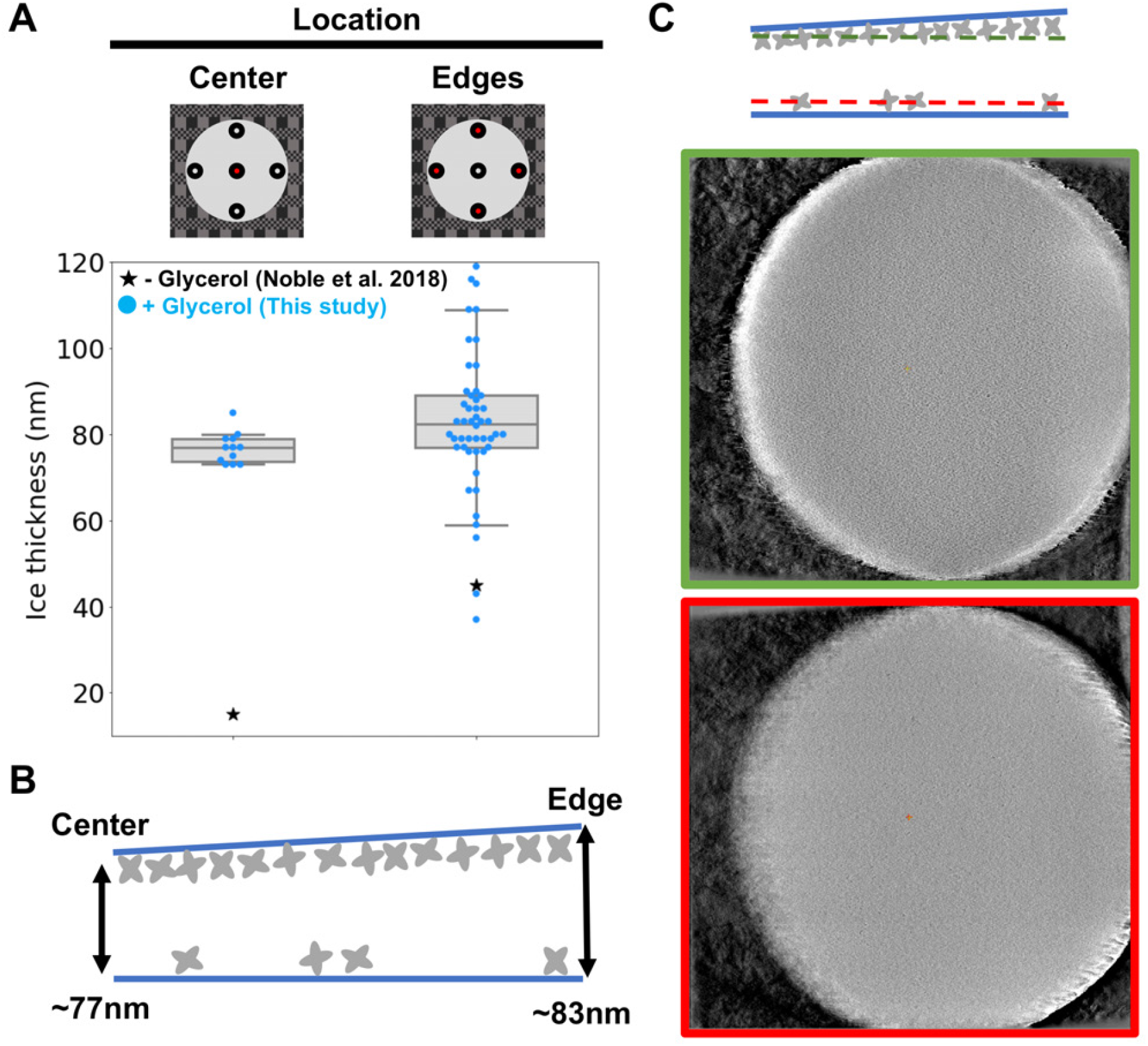
Ice thickness measurements obtained by electron tomography on an aldolase sample in 20% glycerol buffer. **A**. The hole diagrams on the top show the approximate locations of measurements in red, done either in the center or four edge spots. Data points are shown as colored dots (horizontally stacked only to aid clarity) overlaid on a grey box that spans the two central quartiles and is crossed by a vertical line at the median. The whiskers span the range of data between the 5th and 95th percentiles. Black stars represent data from previous studies for samples without glycerol (Rice *et al*., 2018). **B**. Diagram of an average ice cross section for aldolase samples with 20% glycerol integrating all gathered information. The labeled values are the average ice thickness at the edges and center. **C**. Tomogram slices at the height of each of the two particle layers, with border colored according to the lines depicted above in the ice section cartoon.

**Supplementary Figure 13.**
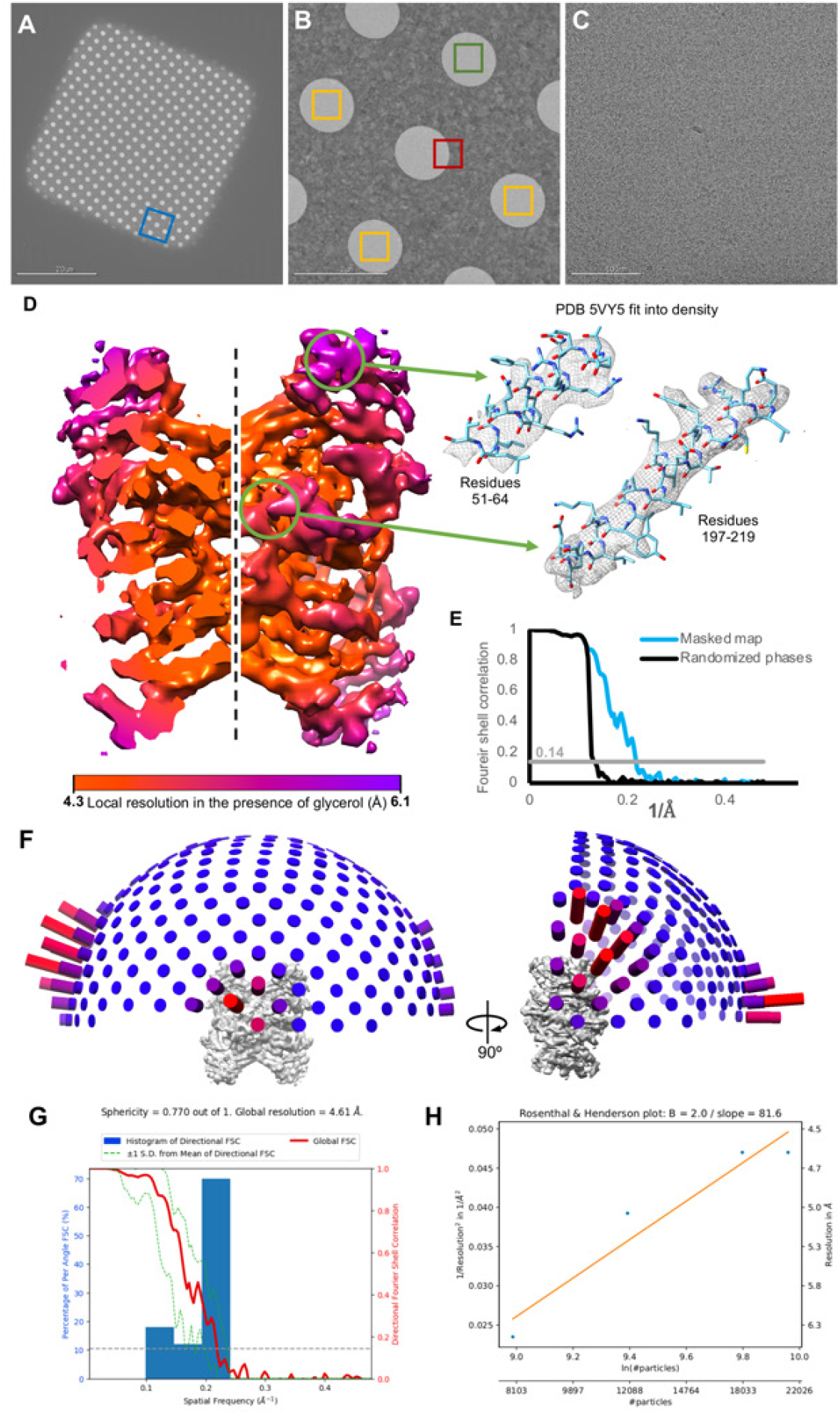
Exemplar micrographs and additional final reconstruction metrics for aldolase imaged at 300KeV in 20% glycerol buffer. **A**. Exemplar grid square, a blue frame marks where micrograph (B) was acquired. **B**. An exemplar “hole” magnification micrograph, with a red square showing where real focus estimation was done, and yellow squares showing where high-magnification micrographs were acquired for single particle analysis. A green square shows where micrograph (C) was taken. **C**. An exemplar micrograph used for single particle picking and analysis. **D**. Reconstruction volume colored by local resolution, as calculates by RELION 3.1 (Zivanov *et al*., 2018). Arrows point at two helices from areas with different overall resolution, with the 5VY5 PDB model fit into the density and displayed inside it. **E**. Fourier shell correlation curves of the masked map and the map with randomized phases, the grey line denotes the 0.143 correlation value. **F**. Angular distribution of views for the final reconstruction done assuming D2 symmetry.**G**. Directional FSC curves showing how resolution varies by orientation (Tan *et al*., 2017). **H**. B-factor plot showing how reconstruction resolution improves as larger subsets of the final particle set are used for 3D auto-refinement (Rosenthal & Henderson, 2003), generated using the bfactor plot.py script provided together with RELION 3.1 source code.

**Supplementary Figure 14.**
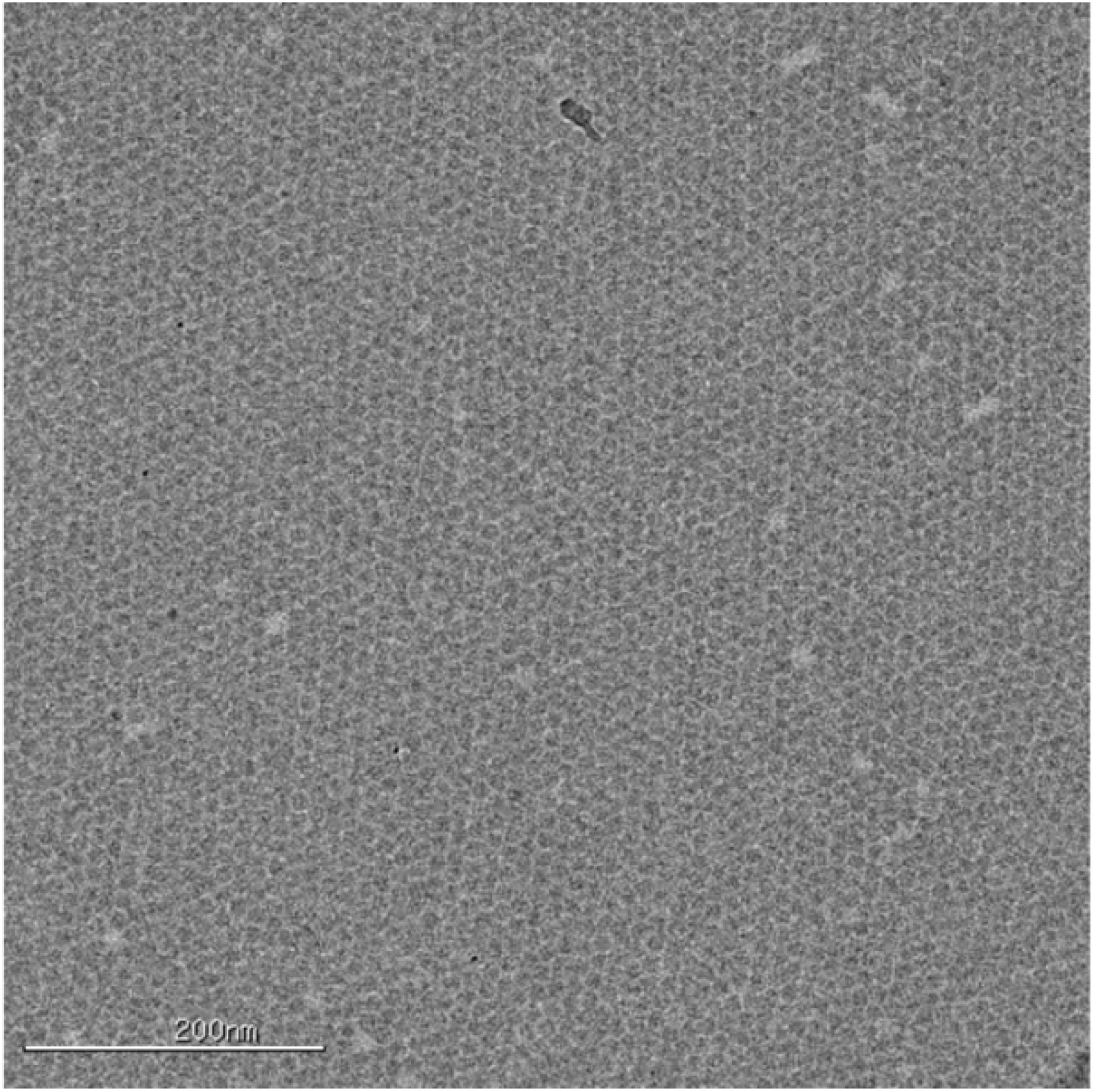
Exemplar micrograph obtained by plunging directly in liquid nitrogen. A clear array of particles, as usually formed in high-concentration samples of apoferritin, is visible in this image taken from a vitrified sample flash frozen in liquid nitrogen after side blotting, as done for negative stain. Image acquired on a Tecnai F20 with a CCD at a magnification corresponding to a pixel size of 1.77Å, and at a defocus -2.5µm.

**Table S1.** Cryo-EM data collection, processing parameters, and statistics.

| EMDB ID | Apo ferritin<br>1.66% glycerol, 200kV<br>EMD-24795 | Apo ferritin<br>20% glycerol, 200kV<br>EMD-24796 | Aldolase<br>20% glycerol, 200kV<br>EMD-24798 | Apo ferritin<br>20% glycerol, 300 kV<br>EMD-24797 | Aldolase<br>20% glycerol, 300kV<br>EMD-24799 |
| --- | --- | --- | --- | --- | --- |
| **Data Collection** |  |  |  |  |  |
| Microscope | Talos Arctica | Talos Arctica | Talos Arctica | Titan Krios | Titan Krios |
| Camera | Gatan K2 Summit | Gatan K2 Summit | Gatan K2 Summit | Gatan K2 Summit | Gatan K2 Summit |
| Voltage (kV) | 200 | 200 | 200 | 300 | 300 |
| Magnification (nominal/at detector) | 36,000/43,478 | 36,000/43,478 | 45,000/54,945 | 29,000/48,732 | 130,000/47,846 |
| Data acquisition software | Leginon | Leginon | Leginon | Leginon | Leginon |
| Exposure navigation | image shift to 16 | image shift to 16 | image shift to 4 | image shift to 16 | image shift to 5 |
| Total electron exposure (e<sup>-</sup>/Å<sup>2</sup>) | 50 | 50 | 80 | 50 | 52 |
| Exposure rate (e<sup>-</sup>/pixel/sec) | 6.0 | 5.8 | 3.5 | 8.1 | 5.7 |
| Frame length (ms) | 100 | 100 | 200 | 100 | 200 |
| Number of frames per movie | 110 | 110 | 96 | 65 | 50 |
| Pixel size (Å/pixel)* | 1.15 | 1.15 | 0.91 | 1.026 | 1.045 |
| Defocus range (μm) | 0.8 to 1.7 | 0.8 to 1.7 | 0.8 to 1.7 | 0.8 to 1.0 to 2.8 |  |
| Micrographs collected (no.) | 1,707 | 2,254 | 741 | 1,859 | 2,591 |
| **Reconstruction** |  |  |  |  |  |
| Image processing package | RELION 3.1 | RELION 3.1 | RELION 3.1 | RELION 3.1 | RELION 3.1 |
| Total extracted picks (no.) | 876,503 | 830,042 | 323,571 | 1,757,706 | 1,255,598 |
| Particles used for 3D (no.) | 875,694 | 829,923 | 142,300 | 778,762 | 143,234 |
| Final Particles (no.) | 598,660 | 683,154 | 28,588 | 778,762 | 21,155 |
| Symmetry | Octahedral | Octahedral | D2 | Octahedral | D2 |
| Resolution (global) (Å) |  |  |  |  |  |
| FSC 0.5 (unmasked / masked) | 2.5 / 2.4 | 2.5 / 2.4 | 6.1 / 3.8 | 2.3 / 2.2 | 7.2 / 6.2 |
| FSC 0.143 (unmasked / masked) | 2.4 / 2.3 | 2.3 / 2.3 | 3.8 / 3.3 | 2.1 / 2.1 | 6.2 / 4.6 |
| Local resolution range (Å) | 2.3 - 2.4 | 2.3 - 2.4 | 3.3 - 4.6 | 2.1 to 2.3 | 4.3 to 6.1 |
| 3DFSC Sphericity (%) | 99.7 | 99.3 | 84.1 | 99.4 | 77.0 |
| Accuracy of translations / rotations | 0.23 Å / 0.236° | 0.23 Å / 0.368° | 0.54 Å / 1.407° | 0.28 Å / 0.406° | 2.02 / 6.82° |
| Map sharpening B-factor (Å<sup>2</sup>) | -19.0 | -36.1 | -56.6 | -117.6 | -197.1 |

## Notes

### Competing Interest Statement

The authors have declared no competing interest.

